# The recombination efficiency of the bacterial integron depends on the mechanical stability of the synaptic complex

**DOI:** 10.1101/2024.04.09.588808

**Authors:** Ekaterina Vorobevskaia, Céline Loot, Didier Mazel, Michael Schlierf

## Abstract

The predominant tool for adaptation in Gram-negative bacteria is a genetic system called integron. Under conditions of stress, it rearranges gene cassettes, ensuring their sampling through expression, to offer a solution for overcoming the initial stress. Integrons are a major actor of multiple antibiotic resistances, a recognized major global health threat. Cassettes are recombined by a unique recombination process involving a tyrosine recombinase – the IntI integrase – and folded single-stranded DNA hairpins – the *attC* sites which terminate each cassette. Four recombinases and two *attC* sites form a macromolecular synaptic complex, which is key to the recombination process and the focus of our study. The bottom strand of all *attC* sites shows highest recombination efficiency *in vivo* than the top one, however, the efficiency still varies several orders of magnitude and the underlying reason remains unclear. Here, we established an optical tweezers force-spectroscopy assay that allows us to probe the synaptic complex stability. We found for seven combinations of *attC* sites great variability in the mechanical stability. Two protein variants also showed a strong influence on the mechanical stability. We then determined the *in vivo* recombination efficiencies of the different *attC* site combinations and protein variants and discovered a strong correlation between recombination efficiency and mechanical stability of the synaptic complex, indicating a regulatory mechanism from the DNA sequence to the macromolecular complex stability. Taking into account known forces during DNA metabolism, we suggest that the variation of the *in vivo* recombination efficiency is mediated strongly by the synaptic complex stability.

## Introduction

Bacterial integrons are the predominant adaptive system among Gram-negative bacteria (*1, 2*). They are found as chromosomal integrons (*3, 4*) and as mobile integrons (*5, 6*). Mobile integrons are part of the horizontal gene transfer and thus one of the three major mechanisms of antibiotic multi-resistance transmittance (*1, 7*). Integrons are genetic elements able to capture and rearrange gene cassettes using site-specific recombination (*2*). Any integron system consists of two major functional parts (*8*): 1) a stable platform, with an integrase gene *intI* and its own promoter (*P*_int_) typically under the control of the SOS response protein LexA (*9*); a cassette promoter (*P*_c_) and the adjacent primary integration sequence *attI*, and 2) a cassette library platform, with promoterless genes stacked one after another, flanked by DNA sequences with imperfect inverted repeat sequences called *attC* sites (Fig. 1A). *attC* sites display high sequence and size diversity, ranging from ∼60 to ∼150 nucleotides (nts), though they are recognized and recombined by the integrase due to a set of characteristics (*10, 11*). All *attC* sites are comprised of a (partial) palindromic sequence, to be able to form a hairpin-like structure (*10*). Once folded as a hairpin, the *attC* sites consist of two integrase binding sites in opposite orientations, the “L-box” and the “R-box” that are separated by an unpaired central spacer (UCS), two to three extrahelical bases (EHBs) and a variable terminal structure (VTS) of varying size (Fig. 1B), structure and yet unknown function (*12, 13*).

**Figure 1:**
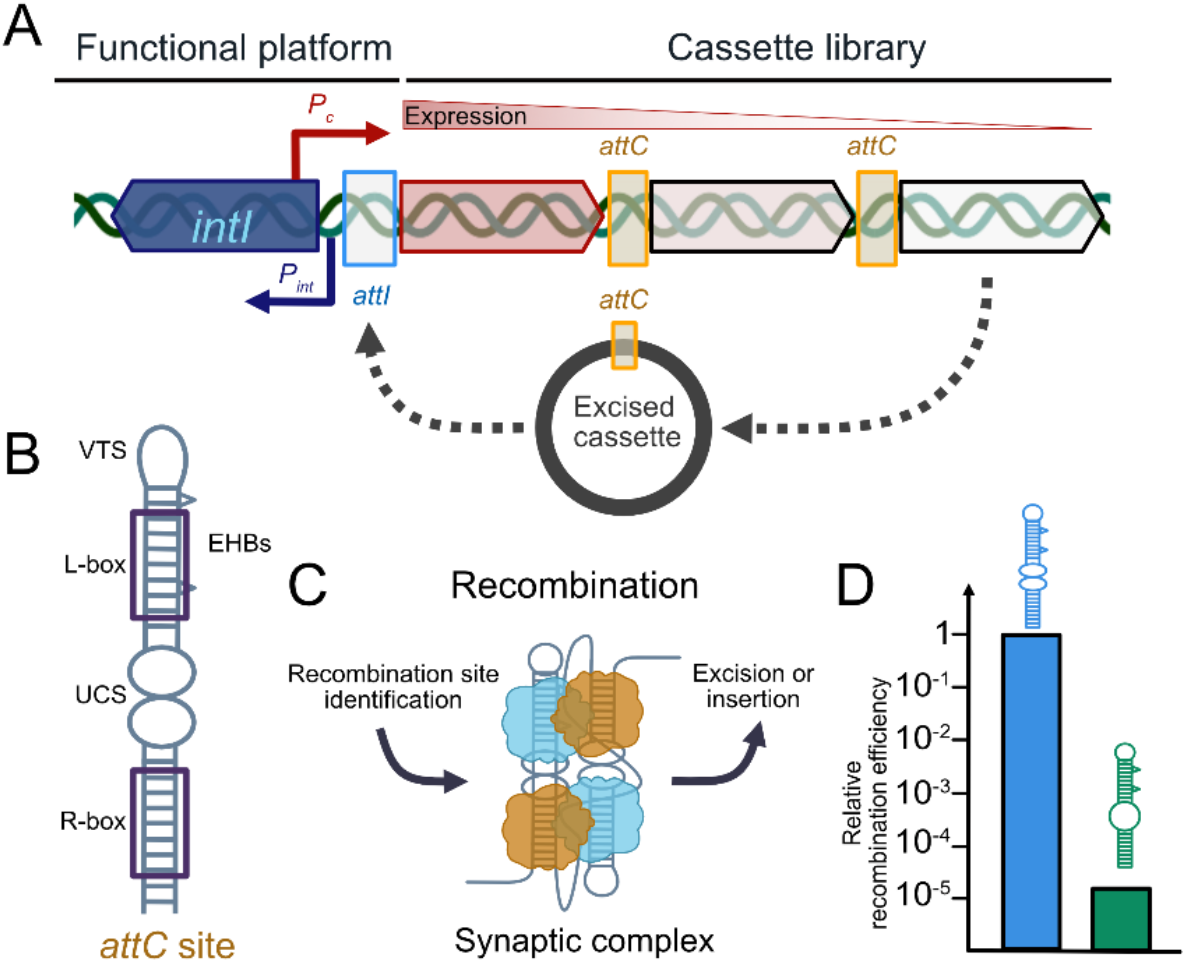
Integron genetic system. **(A)** Schematic representation of the integron: the functional platform contains the integrase gene *intI*, integrase promoter *P*_*int*_, the common promoter for cassettes *P*_*c*_, and the primary integration *attI* site. A cassette library is shown with the expression gradient, depending on the proximity to the *P*_*c*_; *attC* sites are represented with yellow frames. An excised cassette with the putative pathway towards reintegration and adaptation is depicted in grey. **(B)** Sketch of a prototypic *attC* site: extrahelical bases (EHBs), unpaired central spacer (UCS), and variable terminal structure (VTS) are indicated as well as the integrase binding boxes (R-box and L-box). **(C)** Schematic depiction of the role of the synaptic complex in bacterial recombination. Integrase monomers are shown in blue and orange. **(D)** Relative recombination efficiency differences between the integration of two *attC* sites of similar size *in vivo* (*24*).

During SOS response (*9, 14, 15*), the promoters *P*_int_ and *P*_c_ are accessible, thus *intI* and cassettes close to *P*_c_ can be transcribed (*16*). However, most cassettes remain silent storage units until they take part in a recombination process driven by the protein IntI (*17*). IntI recognizes two flanking *attC* sites to excise cassettes from the cassette array and re-integrates excised cassettes mostly at the primary integration site *attI*, allowing its subsequent expression (*2*) (Fig. 1C). Unlike other DNA recombination systems, IntI recombines *attC* sites as single-stranded DNA (ssDNA) elements and forms a unique complex of four recombinases and two folded hairpin structures, that allow the integrase to excise a gene cassette (*18*). The multi-component structure is called the synaptic complex and is crucial for the recombination process. In a crystallographic structure of the synaptic complex of IntI4 with *attC* sites, it was discovered that the EHBs directly interact with specific residues of the integrase, likely mediating allosteric conformational changes upon binding, which result in only two active IntI monomers capable of DNA cleavage in the conserved 5’-AAC-3’ region of the R-box (*18*).

Moreover, integrase shows a strong preference for the bottom strand of the *attC* site, directed by the hairpin structural elements and biological imperative (*19, 20*). This strand preference is crucial to integron function, as it ensures that the cassette is inserted at the *attI* insertion point in an orientation ensuring its expression by the P_*c*_ promoter. The integration process requires a replication event resolving the atypical Holliday junction structure, so the integrated gene cassette is available for expression (*21*).

Earlier studies revealed that the recombination efficiency of the same integrase IntI showed great variability by up to five orders of magnitude on different *attC* site substrates (*10, 21–23*) (Fig. 1D). Extensive *in vivo* studies have been conducted using both naturally occurring *attC* sites of mobile and chromosomal integron systems as well as putative synthetic *attC*s (*10, 11, 24*). The cause of the variation of the recombination efficiency is still unknown and likely connected to several factors. The first important factor is likely the probability to form a DNA hairpin structure. Earlier studies by Loot and colleagues revealed that modifications of the sequence while maintaining the overall general *attC* architecture still led to different free energies of the DNA hairpin and by extension the DNA cruciform. These differences directly affect the ability to form during regular DNA metabolism (*11*). Recent studies uncovered that *attC* sites are prevented to form by single-stranded binding proteins (*25, 26*), however, energetically favorable sequences close to the apical loop of *attC* sites can promote the formation of a mini-hairpin-like structure. The mini-hairpin likely promotes initial IntI recognition and the possibility to displace SSB, causing *attC* folding (*25*). A second important factor is the structural variation of the *attC* hairpin. Here, earlier studies revealed important bases affecting the resulting architecture of the *attC* sites (*10, 13, 20*). Interestingly, a single-molecule optical tweezers study revealed two *attC*_*aadA7*_ folds, a straight and a kinked structure. The kinked structure does not provide a full second binding site of IntI (L-box). Interestingly, the most recombination-efficient *attC*_*aadA7*_ bottom strand has a structural bias towards a straight conformation, offering both binding boxes. The top strand sequence showed more frequently the kinked structure and was found to be 100-fold less efficient to recombine. By extension, correctly folded *attC* sites allow binding of two integrases and thereon form the core structure of the integron recombination system – the synaptic complex. Therefore, a third important factor is likely encoded in the synaptic complex stability, which can be affected by allosteric modulations encoded in the *attC* fold and protein interactions. We hypothesize that the change in recombination efficiency between different *attC* sites correlates with the mechanical stability of the synaptic complex. Less stable complexes are easier to disrupt by forces acting on DNA during processes like transcription, DNA repair, and replication, while more stable synaptic complexes are more likely to initiate recombination. Distinct protein-protein and protein-DNA interactions have been reported for the crystallographic structure of IntI4, suggesting that these might also be key for the mechanical stability of the complex. A better understanding of all the parameters influencing cassette recombination rate could make it possible to predict the dynamics of integron recombination and, more precisely, the cassettes more inclined to recombine and be expressed.

## Results

### Probing the synaptic complex stability using optical tweezers

Here, we studied the mechanical stability of the synaptic complex using optical tweezers. Optical tweezers allow to manipulate single molecules tethered between two microbeads trapped in a focused laser beam to determine structural and mechanical properties with high precision (*27, 28*). The synaptic complex is formed by a pair of *attC* sites of 61 to 131 nucleotides combined with a 110 nucleotides long spacer to mimic an ORF sequence and additional 48 nucleotides for the optical tweezers construct assembly, leading to a ssDNA length of 280 to 420 nt length (Fig. S1). To produce this specific ssDNA sequence, termed double-*attC*, we developed a protocol based on PCR and directed exonuclease activity (see *Materials and Methods*). To prevent DNA cleavage by IntI1 during our experiments, we have decided to use an IntI1 mutation, which substitutes the attacking tyrosine with a phenylalanine (IntI1^Y312F^), termed thereafter IntI1, without impaired binding to *attC* sites (*29, 30*). Such a mutation allows us to characterize binding to ssDNA and, by extension, synaptic complex formation without degradation of the ssDNA substrate. Using IntI1, we have verified double-*attC* binding with electrophoretic mobility shift assays (EMSA, Fig. 3A, Fig. S2). For subsequent optical tweezers experiments, we attached the double-*attC* construct between two 2.5 kbp long double-stranded DNA (dsDNA) handles, carrying either a biotin or a triple-digoxigenin modification at the 5’ end, assembling a single-stranded/double-stranded hybrid DNA molecule (Fig. 2A). Using a microfluidic multi-channel chamber, we formed stepwise a tether between functionalized microbeads trapped by a focused laser beam (Fig. 2B). After successful tether formation, we expose the tether to IntI1 and allow the formation of a synaptic complex at low forces F < 0.5 pN (Fig. 2C). We then pull on the synaptic complex by moving one bead further away with a constant velocity of 250 nm/s. The increased force on the complex leads to its disassembly, eventually, and a distinct length increase (from state 1 to state 2) of the tether, a molecular fingerprint of synaptic complex disassembly. We designed the molecular fingerprint to be 71 nm contour length change. Further bead separation results in increasing forces on the IntI1 bound hairpins, leading to IntI1 unbinding in concert with hairpin unfolding. We then reverse the movement and relax the tether to low forces and allow a new synaptic complex formation. We typically go through tens to hundreds of cycles for individual tethers to acquire many data points for statistical analysis. Experimental data is plotted as a force-extension curve (Fig. 2D), allowing us to identify synaptic complex disassembly by our designed molecular fingerprint (contour length change between state 1 and state 2). The contour length change reflects the transition between two stable states, a formed synaptic complex and a stretched hairpin-bound state, releasing the ssDNA spacer between both *attC* sites (Fig. S3). After identification, we extract structural change information simultaneously with the characteristic disassembly forces and pathways. By exchanging the sequence of the hairpins in the hybrid construct, we investigated the effects of *attC* sequences on the synaptic complex stability.

**Figure 2:**
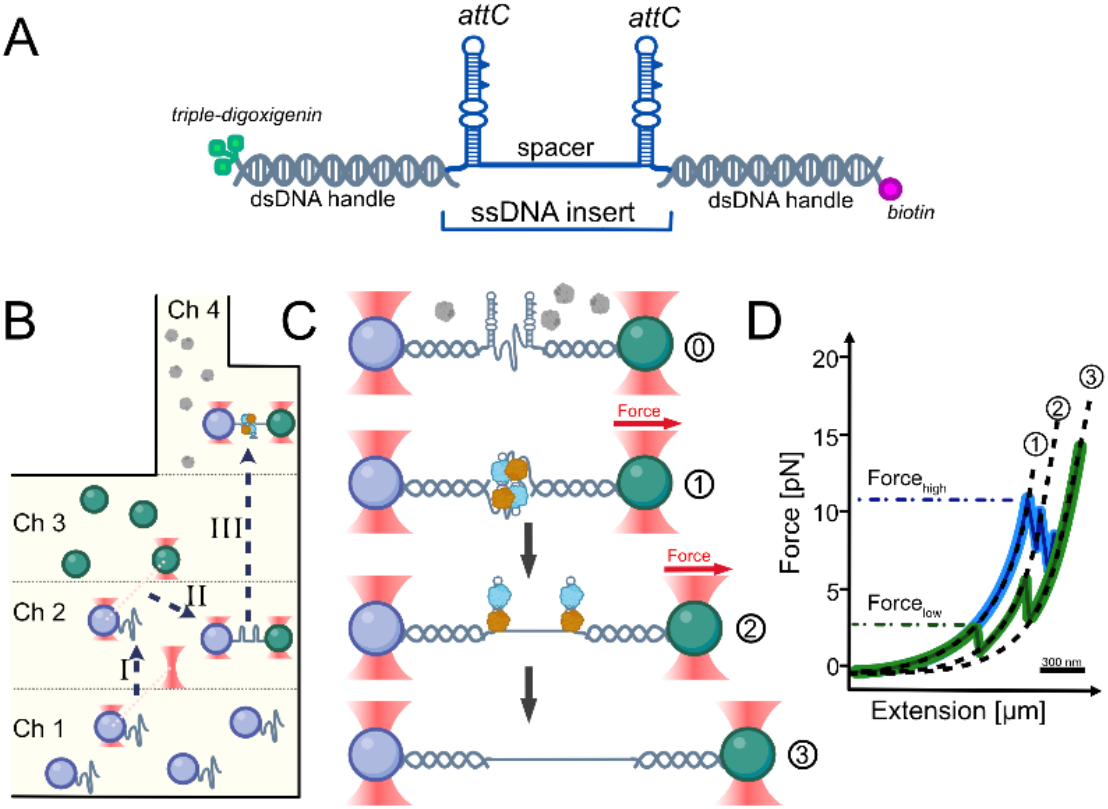
An optical tweezers assay to probe synaptic complex stability. **(A)** Hybrid DNA construct for *in vitro* synaptic complex reconstitution: two *attC* hairpins, a ssDNA spacer for gene cassette imitation, dsDNA handles with 5’ modifications carrying a biotin and a triple-digoxigenin for specific bead attachment. **(B)** Schematic depiction of the experimental procedure. (I) Trapping the anti-digoxigenin bead with the hybrid DNA in Channel 1 and moving to channel 2 containing measurement buffer. (II) Catching the streptavidin bead with the second trap in Channel 3 and moving to buffer Channel 2. (III) Trap calibration, establishing a tether and single tether specificity assessment. Moving to channel 4 for synaptic complex formation in buffer containing IntI1 and synaptic complex stability measurements. **(C)** The tether is exposed to protein monomers in solution (0). By moving Trap 2 away from Trap 1 the tether is stretched and the synaptic complex (1) is disassembled to the hairpin-bound state (2) followed by protein dissociation and hairpin unfolding (3). **(D)** Schematic force-extension curves for high (in blue) and low (in green) recombination efficiency *attC* sites. Synaptic complex disassembly is identified using an established molecular fingerprint ΔL_1-2_=71 nm, and the characteristic force of the event is recorded using the breaking point on the force-extension curve (Force_high/low_). Subsequent events of hairpins unfolding produce the theoretical ΔL_2-3_=82 nm for short *attC* sites or 174 nm for long *attC* sites.

**Figure 3:**
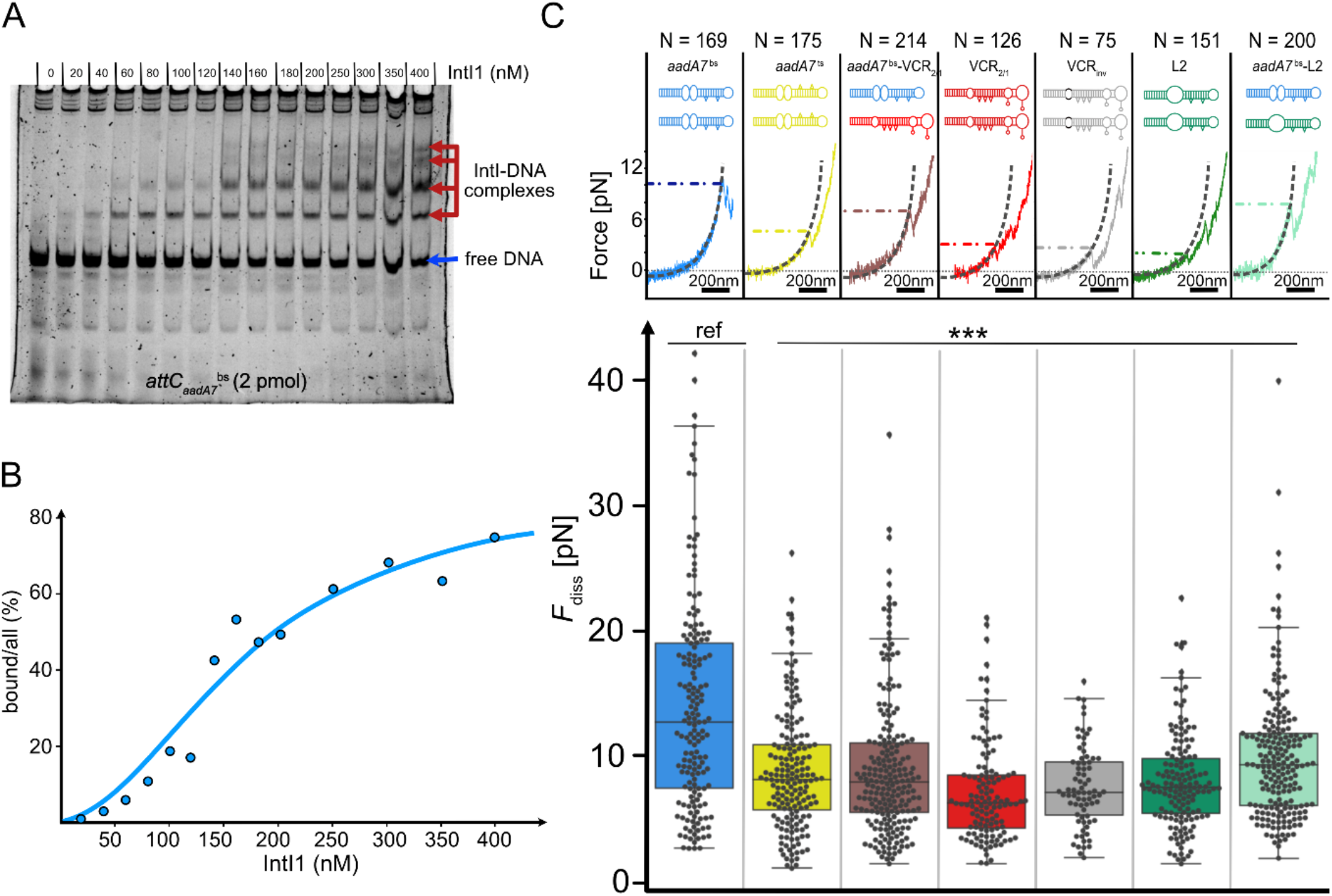
Integrase binding and mechanical stability of the synaptic complex depending on the attC site. **(A)** EMSA with 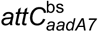 produced up to four shifted bands on the gel. **(B)** Gel data analysis with a ratio of band intensity corresponding to the % of bound DNA. Bound DNA data fitted with the Hill-Langmuir equation. **(C)** Boxplot presents the distribution of synaptic complex disassembly force *F*_diss_ as a measure of mechanical stability for a synaptic complex assembled by integrase IntI1 and different *attC* substrates. Reference 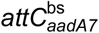 construct is indicated with “ref”, significance is shown with stars (*** for P-value < 0.001, pairwise T-test with Holm correction). The number of measured synaptic complex disassembly events N is given for each *attC* site above the force-extension curve of experimental data with indicated characteristic force.

### Synaptic complexes of different attC sites demonstrate variable mechanical stability

To test the hypothesis if the synaptic complex stability correlates with recombination efficiency, we decided to choose a variety of *attC* sites, which were previously studied for their ability to recombine, and which structural features affect their recombination efficiency. Five *attC* sequences were selected to represent a wide range of recombination efficiencies varying by four orders of magnitude (Table S1). While the general features were overall conserved, the structures still differed locally, as well as in the large VTS, especially for the longer *attC* sites (Fig. S4). We chose both, naturally occurring (originating from mobile *attC*_*aadA7*_ – as well as chromosomal – *attC*_VCR2/1_ – integrons) and synthetically engineered or mutated *attC* sites known for their low recombinogenic properties (*attC*_L2_ and *attC*_VCRinv_, (*10, 14, 24*)). We also investigated one top and bottom strand pair of the same *attC* site (*aadA7*).

All double-*attC* insert sequences were tested for integrase recognition and binding using an EMSA. In brief, purified integrase IntI1 was titrated to the double-*attC* constructs of 286 nt, 280 nt and 420 nt long for *attC*_*aadA7*_, *attC*_L2,_ and *attC*_VCR_ sites, respectively. To ensure stable DNA hairpins and minimize ssDNA during the EMSA, we stapled the hairpins using long complementary oligonucleotide sequences (Fig. S5) and visualized the DNA shift upon protein binding (Fig. 3A, Fig. S2). We observed up to four shifted bands representing all expected molecular complexes formed likely monomer-by-monomer as the integrase concentration was increased. The largest molecular complex with four integrase subunits bound to two *attC* hairpins shows that the reconstitution of the synaptic complex *in vitro* using our designed double-*attC* insert is in principle possible. Using Hill-Langmuir equation (see *Materials and Methods*) we determined the half concentration *c*_1/2_ of IntI1 binding to the most recombinogenic 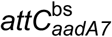 site to be 153 nM ± 22 nM (Fig. 3B). For the top strand 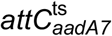 that is widely acknowledged to be a negative control of the integrase binding (*13, 19, 25*), we did observe indeed poor binding on the EMSA with only a monomer band of 7% bound fraction at 350 nM IntI1. The other *attC* sites showed all binding, with an unexpectedly high affinity of IntI1 to *attC*_VCRinv_ (*c*_1/2_ = 135 nM ± 24 nM), moderate affinity of *attC*_VCR2/1_ (*c*_1/2_ = 353 nM ± 64 nM), and the synthetic *attC*_L2_ (*c*_1/2_ = 357 nM ± 40 nM). Interestingly, the determined affinities do not correlate with the reported *in vivo* recombination efficiencies, indicating that the recombination efficiency might be influenced by more than IntI1 binding to *attC* sites but also higher-order organization of the synaptic complex itself. The complex formation, however, cannot be probed using EMSA, but it is possible to assess using optical tweezers.

In the first set of optical tweezers experiments, we assembled a double-*attC* construct with a pair of 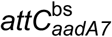 . After tether formation, we incubated the relaxed tether at an IntI1 concentration of 180 nM (in Channel 4; Fig. 2B) to allow for the self-assembly of a synaptic complex. Upon increasing the force, we frequently observed synaptic complex disassembly, showing the designed molecular fingerprint, the contour length increase of 71 nm originating from the release of the ssDNA spacer with bound hairpins and the loss of the synaptic complex assembly (Fig. S3). Synaptic complex disassembly is generally followed by the unfolding of both *attC* hairpins, also indicated by a characteristic contour length increase. We then determined the synaptic complex disassembly forces for each event and found a median disassembly force 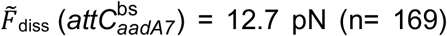 (Fig 3C). The forces were broadly distributed from 5 pN up to 40 pN representing the stochastically driven process of disassembly. Noteworthy, a large fraction of disassembly events occurred at high forces above 25 pN. Testing of the top-strand 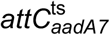 in a subsequent optical tweezers experiment, we observed, to our surprise, frequent synaptic complex formation. We determined a characteristic synaptic complex disassembly force of 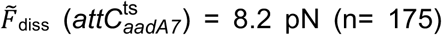, significantly lower than the bottom strand (Fig. 3C). Notably, synaptic complex formation for 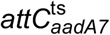 was unexpected, since we observed only low binding affinity of IntI1 using EMSA (Fig. S2).

We repeated these experiments with a double-*attC* construct containing two large VCR_2/1_ *attC* sites and observed a significantly lower synaptic complex disassembly force 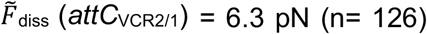 with a narrow distribution (Fig 2C). Both, the mutated *attC*_VCRinv_ and the synthetic *attC*_L2_ sites showed median disassembly forces of 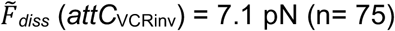 and 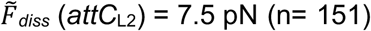. In summary, the double-*attC* constructs revealed a wide spread of median disassembly forces, with 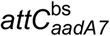 showing the highest mechanical stability. Even though we observe a distinct destabilization of the synaptic complex formed with lower recombination efficient *attC* sites, destabilization does not scale linearly with the disassembly forces.

An important factor that influences integrase binding is correct hairpin folding. It was previously shown that folding of the *attC* influences the recombination efficiency *in vivo* (*11*) and longer *attC* sites (for instance VCR variants with large VTS segments) have more possible folding structures thus reducing the probability of the straight canonical fold. Most probable changes happen at the L-box, located higher on the hairpin stem, closer to the variable terminal sequence. An incomplete binding box folding can either fully prevent integrase from binding or lead to a partially bound conformation. This will either prevent synaptic complex assembly or create a trimer/kinked tetramer assembly that would still be recognized as a synaptic complex in our optical tweezers assay but is expected to have reduced stability as some of the necessary stabilizing interactions can be lost. This impact of overall *attC* length, which creates more misfolded possibilities for the hairpin might be one of the reasons for very low mechanical stability that was demonstrated by the synaptic complex assembled of two *attC*_VCR2/1_ sites. As this site has shown moderate recombination efficiency *in vivo*, a higher stability was expected, based on the overall trend we observe for the *attC* sites with respect to recombination efficiency values (Fig. 3C).

Apart from the correct folding, the sequence of some important structural features was shown to be decisive in rescuing recombination efficiency. As demonstrated by (*13*) only three nucleotides change in the unpaired central spacer of the hairpin can already invert preferred *attC* conformation and increase recombination efficiency. Inverting the nucleotides in the unpaired central spacer reduces the efficiency of the *attC*_VCRinv_ three orders of magnitude compared to the *attC*_VCR2/1_ (*10, 23*), however, there is no noticeable change in synaptic complex mechanical stability. This might indicate that the folding geometry plays a more important role in the synaptic complex stability than the sequence itself. Notably, in an *in vivo* context and precisely in Mobile Integrons, rarely two identical *attC* sites take part in a synaptic complex, but typically two *attC* sites of different sequence and structure are involved. Therefore, we constructed two heterogenous double-*attC* tethers, one comprised of 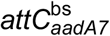 and *attC*_VCR2/1_ and another of 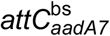 and *attC*_L2_, because both, *attC*_VCR2/1_ and *attC*_L2_, showed the lowest mechanical stability in the homogenous double-*attC* tethers. We next probed, if a synaptic complex formed with a hetero-double-*attC* tether containing *aadA7* ^bs^ can increase the overall stability. Indeed, combining a “weak” *attC* site with a “strong” *attC* site partially rescued the mechanical stability of the synaptic complex with 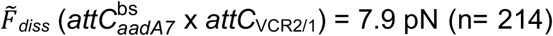 and 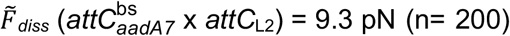 (Fig. 3C). This finding emphasizes the intricate allosteric control of the synaptic complex through the DNA substrate.

### Point mutations in the integrase modulate synaptic complex stability

To investigate the role of the integrase on synaptic complex stability, we focused on two major features: protein-DNA interactions to the adjacent *attC* hairpin and protein-protein interactions within the tetramer. Both are considered to be involved in complex stabilization (*18*). To investigate the role of protein-DNA interaction we aimed at an integron-specific I2 α-helix of the attacking subunit that interacts with the adjacent *attC* in the region of the second extra helical base *in trans* (*31*) (Fig S6A). We designed an alanine mutant that has two conserved residues, lysine and tyrosine, mutated to alanines – IntI1^Y312F-K219A-Y220A^ (from now on IntI1^A^). The subunits in a tetramer interact through C-terminal chains with a small α-helix that is buried in the neighboring subunit, strengthening the synaptic complex (Fig S6B). To test how the C-terminal interaction influences complex stability, a truncated mutant IntI1^Y312F-A321X^ (from now on IntI1^ΔC^) was designed that lacked the C-terminal α-helix.

In the context of the synaptic complex, alanine mutations will only affect two subunit interactions out of the tetramer, because only active subunits have been reported (*18, 31*) to show interaction to the adjacent *attC*, therefore we would expect only a mild effect on the synaptic complex stability. In initial experiments, we probed the binding of IntI1^A^ to our double-*attC* construct using EMSA. IntI1^A^ bound stably one to four monomers to the construct with similar behavior as IntI1, showing increased dimers and tetramers at elevated protein concentrations (Fig. S2B). We observed a slightly increased affinity of IntI1^A^ with a *c*_1/2_ (IntI1^A^) = 100 nM ± 14 nM vs *c*_1/2_ (IntI1) = 152 nM. Subsequently, we switched to the optical tweezers assay with the hybrid DNA construct containing two highly recombinogenic 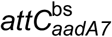 hairpins to probe the synaptic complex stability with IntI1^A^. We observed that despite the mutations, IntI1^A^ formed well-defined synaptic complexes with a moderately reduced median disassembly force of 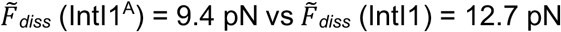 (Fig. 4A). This observation agrees with our hypothesis that likely only two subunits are affected by the mutations and it suggests that the αI2 interaction might be more important for the internal organization of the synaptic complex to determine the catalytic active subunits, but the αI2 interaction has only a minor effect on the overall mechanical stability of the synaptic complex.

**Figure 4:**
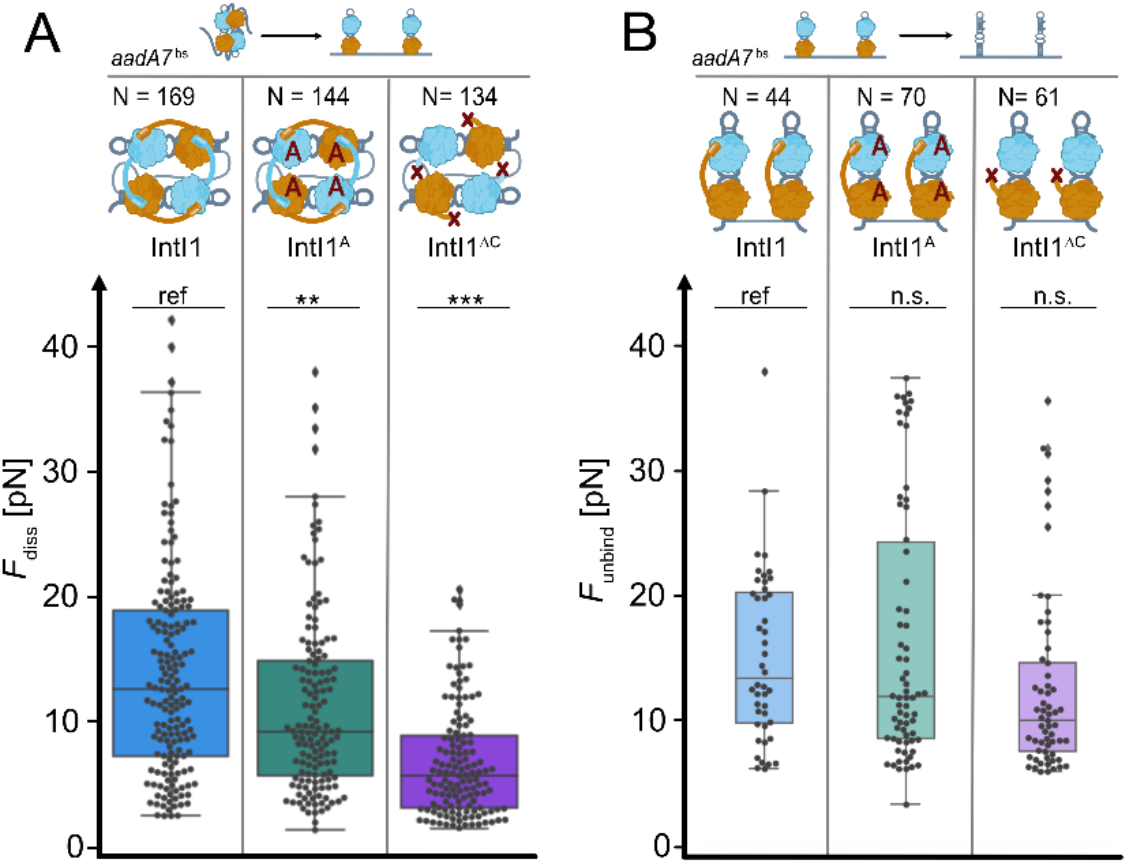
Stability of the synaptic complex depending on protein variants. **(A)** Boxplot presents the distribution of synaptic complex disassembly force as a measure of mechanical stability for a synaptic complex assembled by integrase variants and the most efficient 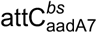 substrate. **(B)** Boxplot presents the distribution of hairpin unbinding force *F*_unbind_ reflecting the strength of integrase binding the *attC* hairpin in case of no previous synaptic complex formation. Reference 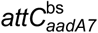 construct is indicated with “ref”, significance is shown with stars (** for P-value < 0.01, *** for P-value < 0.001), “n.s.” – not significant, pairwise T-test with Holm correction. The number of measured synaptic complex disassembly/hairpin unbinding events is given for each protein mutant in N. A catalytically inactive full-length protein IntI1 is colored blue, alanine mutant IntI1^A^ is colored teal, and C-terminal truncated mutant IntI1^ΔC^ colored violet.

For the IntI1^ΔC^ variant, we expect a larger effect on the synaptic complex stability, since it involves direct protein-protein interactions between all four subunits. Using EMSA, we confirmed that IntI1^ΔC^ still binds the double-*attC* construct. However, we observed a lower affinity with *c*_1/2_ = 180 nM ± 17 nM and only a one or two subunit binding was observed on the gel shift (Fig. S2B), even at 600 nM IntI1^ΔC^, in agreement with an earlier study on an IntI1 homolog suggesting that the C-terminal extension might stabilize the tetramer formation (*18*). Looking at the structural organization, such a C-terminal extension might also affect the dimer stabilization on the same *attC*. Upon closer inspection of our EMSA, we observed that IntI1^ΔC^ showed a strong monomer binding, and at elevated concentrations, we observed only two IntI1^ΔC^ monomers binding to our double-*attC* construct. Interestingly, the monomer fraction was not depleted at higher concentrations as observed for IntI1 and IntI1^A^, suggesting that the second band originates from two independent monomer binding events, supporting that the loss of the C-terminal tail could destabilize dimer formation. Unexpectedly, in our optical tweezers assay, we did observe distinct synaptic complexes formed by IntI1^ΔC^ with our hybrid-double-*attC* construct. Yet, IntI1^ΔC^ synaptic complexes showed a pronounced more than 2-fold reduction in median disassembly force 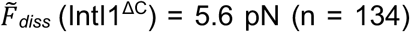 compared to IntI1. Interestingly, the observed disassembly forces were also very narrowly distributed with only a few events exceeding 15 pN (Fig. 4A). These results indicate that while IntI1^ΔC^ is able to form a synaptic complex, it is significantly weakened by the reduction of important protein-protein interactions between all subunits of the complex.

We wondered how the mutations affect the binding strength of IntI1 to the *attC* hairpins themselves. To extract binding strength, we selected force-extension traces that did not show a synaptic complex but started in state 2 (Fig. 2C) showing a well-defined hairpin-bound state due to the distinct contour length fingerprint (see *Materials and Methods*). Using this subset of our data, we determined the median unbinding forces 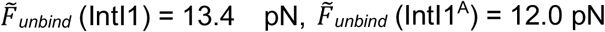, and 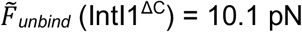, showing only little variation. Applying a pairwise T-test with Holm correction (*32*), we did not observe a significant difference in *F*_unbind_ between the mutants and IntI1. Noteworthy, from the distribution of unbinding forces, it appears that IntI1^ΔC^ had an overall more compacted distribution with only a few outliers at higher forces. While we cannot distinguish monomer and dimer unbinding in our data, we speculate that the higher force events might originate from stable dimers, which would be impaired by the C-terminal α-helix deletion - in agreement with our EMSA observations.

### Classification of attC-IntI interactions reveals a non-canonical disassembly pathway

From a cycle of our single-molecule force spectroscopy experiment, we obtain also structural information on whether and what kind of protein-DNA interactions formed. Within our data, we classified three categories of specific single-molecule events, depending on the mode of integrase interaction with the DNA as well as the pathway that follows a synaptic complex disassembly event. We defined the following three categories: (i) initial hairpin-bound state, but without a synaptic complex; and synaptic complex, leading (ii) to a non-canonical pathway after disassembly or (iii) to a canonical pathway after disassembly (Fig. 5, Fig. S7). We distinguish between the canonical and the non-canonical pathway depending if the tether follows the expected 71-nm fingerprint in a single step or with additional intermediate steps, respectively. We extracted the occurrences for the three different categories for all our IntI-*attC* site experiments. Notably, for either all *attC* variants with IntI1 or 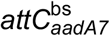 with the IntI1 variants, we observed the hairpin-bound state without a synaptic complex formation (category (i)) with the lowest frequency, with occurrences between 8% for *attC*_L2_×*attC*_L2_ sites and up to 30-33% for IntI1 and variants with 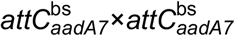 .

**Figure 5:**
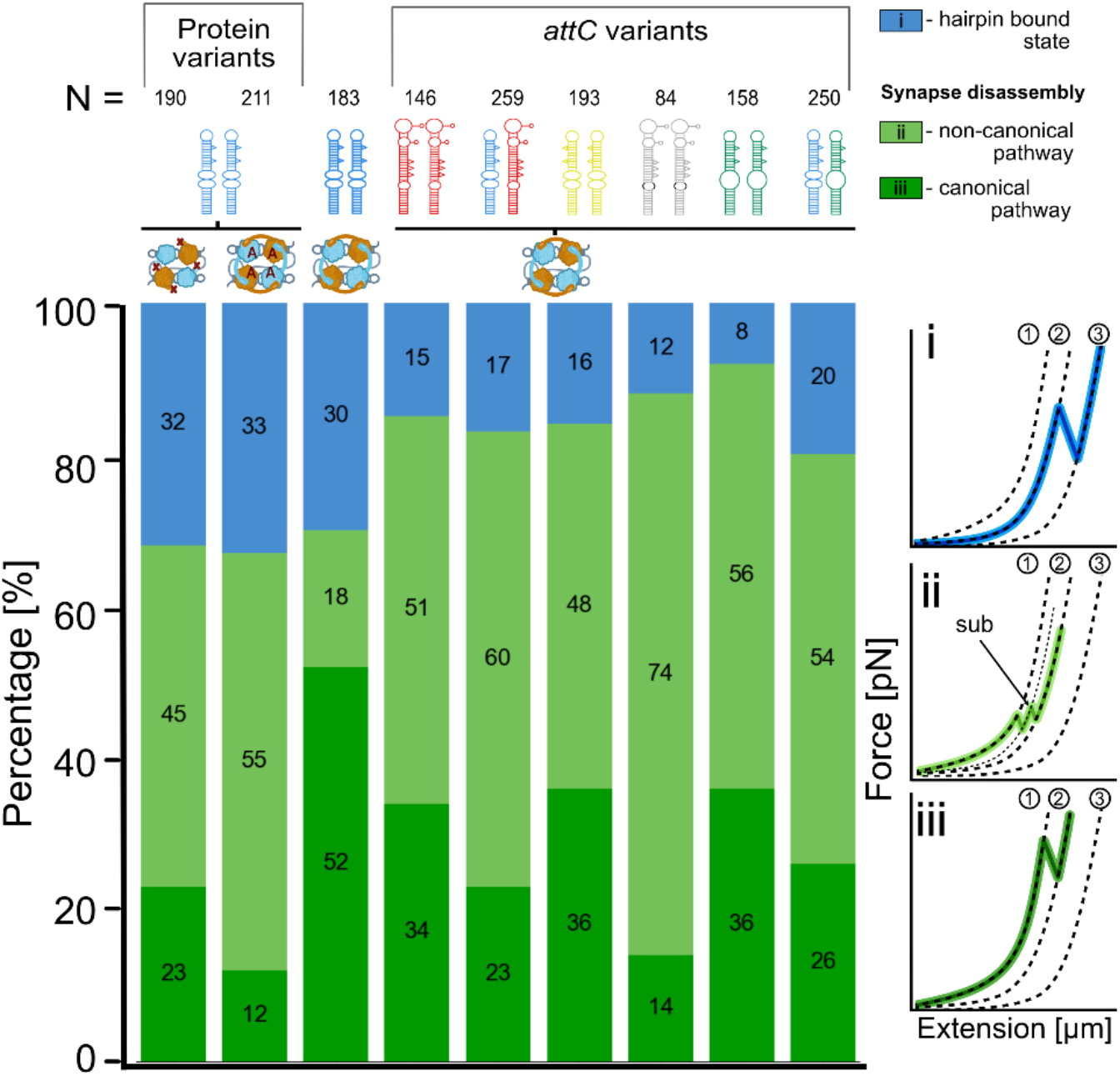
Classification of integrase interaction events with frequency analysis. The stacked chart describes the frequencies of specific integrase interaction events for all protein-DNA combinations. The corresponding event classification is color-coded and given on the right panel. Dashed lines represent the hWLC curves characteristic to the stable states of the tether: (1) – assembled synaptic complex formed by four integrase subunits and two *attC* hairpins; (2) – hairpin bound state with two integrase subunits binding each of the *attC* hairpins separated by a stretched ssDNA spacer; (3) – fully unfolded state with no integrase subunits bound and both hairpins fully unfolded and stretched. A substate (sub) was presented as an intermediate binding step between states (1) and (2). It occurred after the synaptic complex disassembly but before the spacer between the hairpins is fully stretched and might be an unpredicted interaction between a bound hairpin and a spacer. The total number of events is given for each protein-DNA combination in N.

Synaptic complex disassembly was the most frequent integrase interaction event in our experiments (categories (ii) and (iii)) in the presence of IntI1. In approximately 67 to 92% of the cases, we observed a synaptic complex formation (mean ± SD = 80 ± 9%). Based on the design of the double-*attC*, the expected stable state after synaptic complex disassembly constitutes two folded, IntI1-bound *attC* hairpins separated by a stretched ssDNA spacer. Such events are named hereafter a canonical pathway leading to the 71-nm molecular fingerprint achieved in one single step. Such behavior was observed in 12-52% of the traces, with 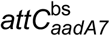 showing the highest occurrence, while the protein variants IntI1^A^ and IntI1^ΔC^ showing the lowest frequency of canonical pathways. However, in some cases upon synaptic complex disassembly, we observed either a distinct substate or an ensemble of short-lived multiple states prior to reaching the stable IntI1-bound *attC* hairpins (Fig. 5, Fig. S7). We called these events non-canonical pathways. Such instances were present for all investigated *attC*-protein combinations and occurred more frequently for mechanically less stable synaptic complexes (either low-efficiency *attC* sites or integrase variants). The mean synaptic complex disassembly force prior to a non-canonical pathway of all investigated IntI:*attC* complexes was 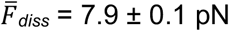 (mean ± SEM). This suggests that low mechanical stability of a synaptic complex allows for the formation of intermediate states during the disassembly pathway. Alternatively, the synaptic complex was incompletely formed, thus less stable, allowing additional intermediate state formation during the disassembly pathway (Fig. S7). Category events were observed in 18 to 74% of the traces, with the more stable 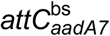 showing the lowest frequency of non-canonical pathways. *attC*_VCRinv_ showed the highest fraction of non-canonical pathways and one of the lowest recombination efficiencies in literature (*10, 23*).

The transient states on the non-canonical pathway have not been observed before and understanding the origin remains challenging. For the case of a single distinct substate, which we observed for different *attC* variants and protein variants, we determined a contour length change of ΔL_1-sub_ = 44.6 ± 3.9 nm (mean ± SD, n = 25). Such a contour length increase is incompatible with a structural change occurring within the synaptic complex (Fig. S8), therefore, we hypothesize that after the synaptic complex disassembly a new interaction between the IntI1-bound hairpins and the single-stranded DNA forms before the spacer is fully stretched.

While the exact structure of this substate remains elusive, our hypothesis suggests that IntI1 also interacts with unstructured ssDNA directly. We tested this hypothesis using correlative force-fluorescence microscopy, where we presented in an optical tweezers assay dsDNA or ssDNA of the same sequence and imaged IntI1 binding behavior. To this extent, we generated a fusion protein of IntI1 with mEGFP and observed to our surprise, frequent and strong binding to ssDNA, even at high tension up to 40 pN (see *Supplementary Information* and Fig. S9). This observation supports our hypothesis that IntI1 can bind the ssDNA spacer of our construct and thus the observed substate can be formed (Fig. S8B).

### Integrase-mediated in vivo recombination efficiency studies

In order to relate the different mechanical stabilities of the synaptic complex to *in vivo* recombination, we measured in an *attC* × *attC* deletion assay the recombination efficiencies. During the deletion assay, a suicide ssDNA is delivered by means of conjugation to recipient strains containing a vector expressing the integrase IntI1^wt^ or variants IntI1^wt-A^ and IntI1^wt-ΔC^ (*10*) (see *Materials and Methods*; Fig. 6A). We determined the excision properties of 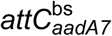 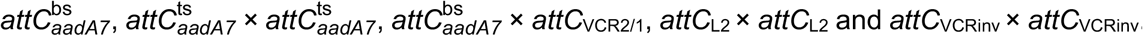. We observed high recombination efficiency of the 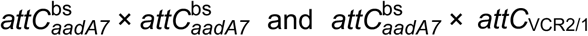 cassettes injecting the bottom strands (8.6×10^-4^ and 4.3×10^-5^, respectively; Fig. 5B). In the absence of IntI1^wt^ we observed much lower recombination efficiencies, indicating that the observed events were integrase-mediated. While injecting the top strand of *aadA7* 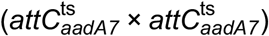, in agreement with earlier reports, we observed a large decrease in the excision efficiency compared to the injected bottom one (to 8.8×10^-6^; Fig 6B). Notably, these recombination events predominantly originate from the newly synthesized bottom strand as previously shown (*10*). We cannot exclude the possibility of top-strand recombination, however, the probability of such events is reportedly low (*10*). Noteworthy, the detected excision efficiency remained integrase-mediated. Both *attC*_L2_ × *attC*_L2_ and *attC*_VCRinv_ × *attC*_VCRinv_ pairs have demonstrated a meager excision efficiency of less than 10^-6^ and did not exceed the background rate obtained in the absence of IntI1^wt^. Background recombination was recently reported to occur due to RecA-independent microhomology recombination (*33, 34*). Thus, observed recombination events close to the detection limit cannot unambiguously be assigned to integrase-mediated recombination, though they were significantly lower than the efficiencies obtained with *attC*_*aadA7*_ and *attC*_VCR2/1_ sites.

**Figure 6:**
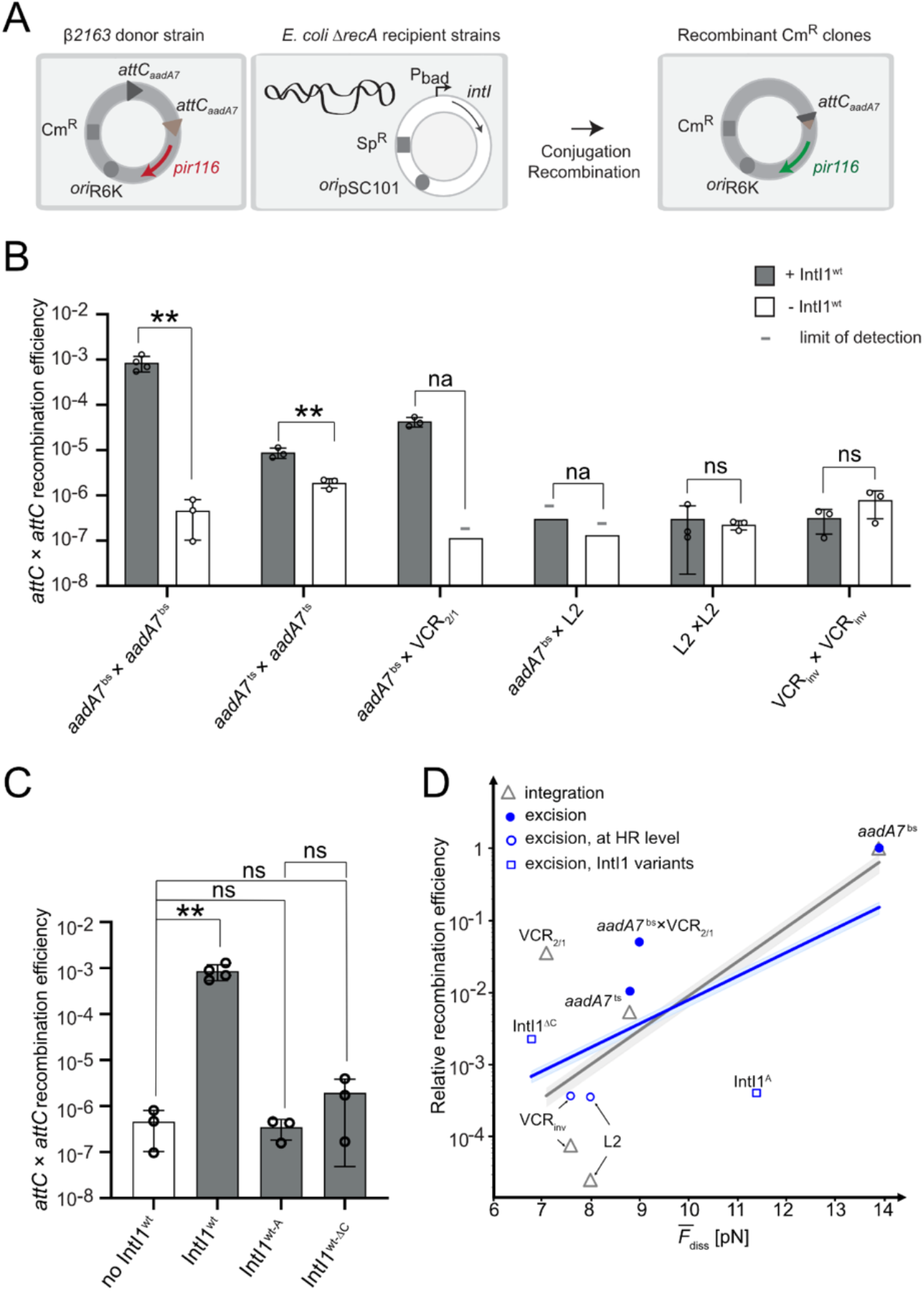
Recombination efficiencies and correlation. **(*A*)** Experimental setup of the cassette excision assay. The pSW23T suicide vector (chloramphenicol resistance Cm^R^) containing the two *attC* sites (grey and brown triangles) is delivered by conjugation from the β2163 *E. coli* donor strains to the MG1655 Δ*recA E. coli* recipient strain. The recipient strains contain a plasmid (spectinomycin resistance Sp^R^) expressing the integron integrase variants under the control of P_BAD_ promotor. pSW23T can only replicate after recombination between both *attC* sites to ensure the expression of the *pir116* gene (green arrow). Recombinant clones are selected on appropriate Cm containing plates. **(*B*)** Cassette excision frequency testing several *attC* sites. Recipient strains containing the pBAD43 integrase expressing vector (+ IntI1^wt^ ) and control strains containing an empty pBAD43 vector (-IntI1^wt^). (-) indicates the recombination frequency was below detection level, indicated by the bar height (limit of detection). Bar charts show the mean of at least three independent experiments (n≥3, individual plots). Error bars show the sd. Statistical comparisons (T-test) are as follow: (na. (not applicable), ns. (not significant), ** P-value < 0.01). **(*C*)** Cassette excision frequency testing several integrase variants. The graph representing the recombination frequencies obtained with two *attC*_*aadA7*_ sites is shown. Bar charts show the mean of at least three independent experiments (n≥3, individual plots) and error bars show the sd. Statistical comparisons (T-test) are as follow: (ns. (not significant), **P-value < 0.01). **(*D*)** Correlation plot for relative recombination efficiency and mean synaptic complex disassembly force showing two sets of data: integration recombination efficiency based on literature data (*10, 11, 13, 23, 24*) (grey triangles) and excision recombination efficiency measured in this study (blue). Different *attC* sites are indicated with circles, IntI1 variants with squares. Filled circles indicate that obtained recombination efficiency was IntI1-mediated, whence empty circles and squares show efficiency at the level of homology recombination.

Interestingly, both of our introduced IntI1 mutations IntI1^wt-A^ and IntI1^wt-ΔC^, have greatly diminished the 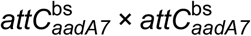 excision efficiency, compared to the IntI1^wt^, differing 3-to-4 orders of magnitude (Fig. 6C). Again, the recombination efficiency of the integrase mutants did not differ from the background homology recombination, suggesting either full loss or strong reduction of activity.

## Discussion

### Synaptic complex of low mechanical stability disassembles at cellular forces

Having determined the *in vivo* recombination efficiencies and the mechanical stabilities of the synaptic complex, we plotted the relative recombination efficiency versus the mean synaptic complex disassembly force (Fig. 6D). We find a clear correlation between both parameters strongly suggesting that the mechanical stability modules the recombination efficiency. Subsequently, we plotted literature values of the relative recombination efficiencies originating from integration (*attC* × *attI*) complexes, which also follow the correlation trend (Fig. 6D, grey).

Even though the excision recombination efficiencies of the tested double-*attC*s range up to 4 orders of magnitude *in vivo*, it does not come to scale in their characteristic disassembly forces. From mechanical stability experiments, it is well known that the rate of disintegration scales exponentially with the applied force, a mechanism which is likely also at play during the synaptic complex disassembly (*35*). Comparing the lower efficiency *attC*s between each other, we observe that 10- and 100-fold differences in recombination efficiency *in vivo* manifest as synaptic stability differences at the scale of only 1 pN. Notably, we did not observe mean disassembly forces below 6 pN, indicating that the system has reached a low threshold of sustainable stability. We propose that the threshold correlates with the average tensile forces that act on the DNA *in vivo* during multiple cellular processes, such as translation, transcription, or replication (*36–39*) (Fig. 7). Less stable complexes dissipate less work prior to disassembly since the surrounding DNA will require less stretching (see *Supplementary Text* and Fig. S10). In consequence, mechanically weak complexes, will have a very short lifetime, due to the exponentially accelerated disassembly kinetics with force, effectively inhibiting successful completion of recombination.

**Figure 7:**
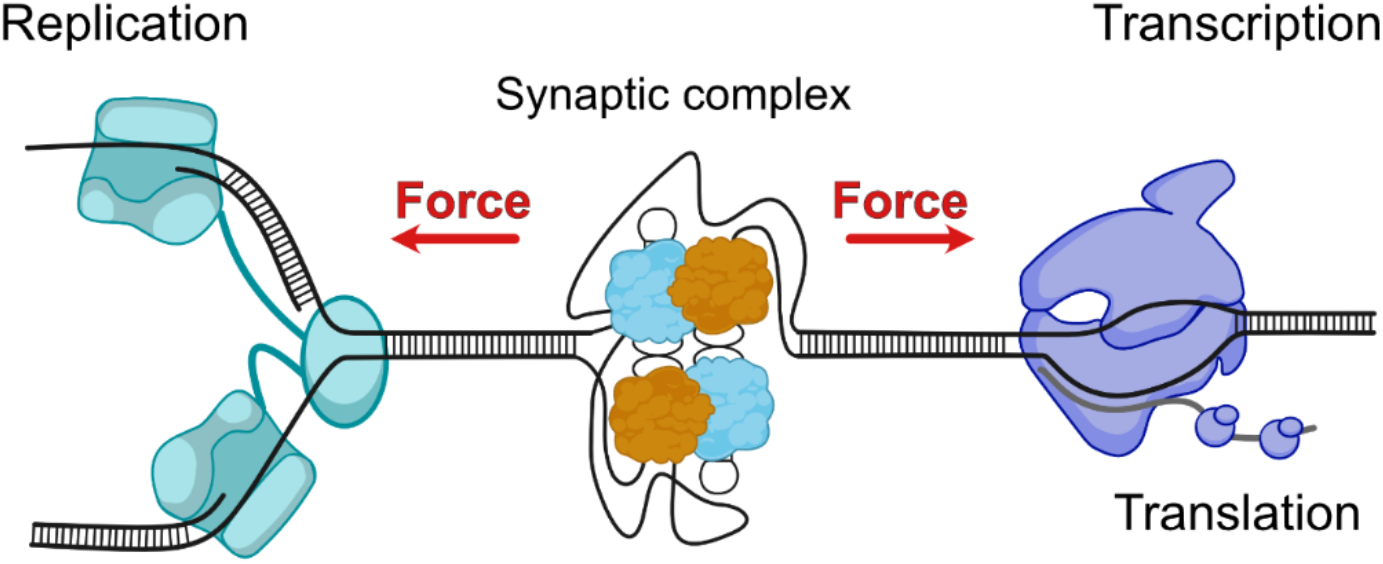
The proposed regulation model. The model of the interplay between the variable synaptic complex stability and the cellular processes that introduce tensile stress on the DNA *in vivo*.

We also discovered a non-canonical pathway after synaptic complex disassembly, which occurred more frequently for low recombination efficient *attC* sites. At the same time, we discovered, that IntI1 strongly, non-specifically binds to ssDNA. Is there an additional benefit of strong ssDNA binding of IntI1? In bacterial cells, the two primary sources of ssDNA are conjugation (*40*) and replication (*41*). Both processes favor integrase-mediated recombination (*2*). Though, it has been shown that recombination efficiency also strongly depends on the supercoiling state of DNA, suggesting that IntI1 recombination occurs with extruded *attC* cruciform-like structures (*11*). Supercoiling is an important regulatory mechanism in bacteria to ensure DNA stability but also chromosome compaction, and typically mediated by topoisomerases, like gyrase (*42, 43*). Yet, the occurrence of transient single-stranded DNA during negative supercoiling is not necessarily leading to immediate cruciform formation, therefore through the non-specific ssDNA binding of IntI1, a continuous supply of monomers close to potential *attC* sites increases the local concentration of the protein, promoting hairpin binding and extrusion. As a consequence, this mechanism likely increases the frequency of synaptic complex assemblies. In the case of a synaptic complex breakdown, the non-specific ssDNA-bound integrases could act as a rescue anchor for the reformation of a synaptic complex.

Recombination efficiency of *attC* sites varies several orders of magnitude and the underlying reason remained unknown. Using optical tweezers, we determined that mechanical stability of synaptic complexes correlates with *in vivo* recombination efficiencies. This correlation illustrates the allosteric modulation of the macromolecular complex stability of the integron by the DNA structure. We anticipate that such a dependency of recombination efficiency on mechanical stability might be at play in numerous recombination processes.

## Materials and Methods

### Double-attC DNA construct

To provide a substrate with different *attC* sites for the synaptic complex reconstitution *in vitro*, we designed a variable ssDNA insert. This insert carries a minimal set of features for the synaptic complex formation: two *attC* hairpins and a spacer that allows antiparallel orientation of the *attC*s when forming a synaptic complex and provides a designed molecular fingerprint identifying synaptic complex disassembly (Fig. S1). In brief, the insert is from 280 nt to 420 nt long, including two *attC* sequences flanked on both sites by short and long spacer sequences as well as overhangs for annealing to dsDNA handles for subsequent optical tweezers experiments. The ssDNA spacer between the *attC*s was designed using Mfold web server (*44*) to avoid additional secondary structures besides the two designed *attC* hairpins. The inserts for the different *attC* sites were synthesized on a plasmid (pUC57-simple; Genscript, USA), full insert sequences are provided (Table S3E). To generate ssDNA inserts, we developed a two-step protocol using selective strand degradation with lambda exonuclease (New England Biolabs, USA) based on (*45*). During amplification of the insert sequence with PCR (primers listed in Table S3B), we introduced on the strand to be degraded a 5’-end phosphorylation, which is recognized by lambda exonuclease for preferred degradation. The resulting ssDNA was used for EMSA as well as for single-molecule optical tweezers force spectroscopy assay.

For hetero-*attC* site constructs generation, we used In-fusion cloning (TakaraBio, Japan) substituting the second *attC* pair in the double-*aadA7* ^bs^ construct for VCR_2/1_ or L2. We used the homo-*attC* construct pair plasmids as templates for the vector and insert of the cloning procedure. The rest of the construct preparation was the same as described above.

### Optical tweezers DNA construct

To avoid surface interactions of the synaptic complex with the beads, we introduced long dsDNA spacers (named DNA handles) between the ssDNA insert and bead surface. 2.5 kbp DNA handles were produced using PCR from λ-DNA (Promega, USA) with a primer carrying either a 5’ biotin or 5’ triple-digoxigenin modification for specific binding to the beads (primers listed in Table S3A) and a 15 bp long sequence carrying double nicking enzyme sites on the reverse primer. The corresponding nicking enzymatic reaction using Nb.BsmI and Nb.BsrDI (New England Biolabs, USA) created the complimentary overhangs to match the insert (*13*). For the final assembly step, a nicked biotin functionalized handle was incubated with the insert for 10 minutes at room temperature to promote initial annealing of the fragments, then mixed up with the nicked triple-digoxigenin handle and put in a thermocycler cooling from 90ºC to 16ºC for coupling (in the ratio 1.2:1:1 for the insert and both handles). It was followed by 2-hour ligation reaction using T4 ligase (Jena Biosciences, Germany) at 16°C to ligate the assembly.

### Single-molecule optical tweezers experiments

All measurements were performed on a commercial, high-resolution correlative fluorescence optical tweezers instrument (C-trap, LUMICKS). As a preparation step the DNA construct was incubated with 2.09 μm anti-digoxigenin coated polystyrene beads (0.1% w/v, Biozol, Germany) for 15 min at room temperature. The construct was attached to the 1.76 μm streptavidin coated polystyrene bead (1% w/v, Spherotech, USA) using laminar flow of the microfluidic device (Fig. 2B). The established tether composed of a single hybrid DNA molecule was confirmed using the real-time force-extension curve analysis by comparing to the expected calculated contour length of the tether (5 kbp dsDNA ≈ 1.7 μm) as well as the steepness of the slope reflecting the persistence length. The traps were calibrated to have a stiffness of 0.4-0.5 pN/nm. Then the tether was moved to channel 4 of the flow chamber that contained integrase protein diluted in measuring buffer to 120-180 nM and the relaxed tether was allowed to form a synaptic complex (Fig. 2B). The measurements were conducted without the flow, mostly in the protein channel with occasional quality checks in protein-free channel. Force-spectroscopy measurements were performed in the microfluidic chamber of the setup in channels 2 and 4, in PBS buffer (Gibco, USA) with the addition of enzymatic POC oxygen scavenger system (*46*) at a constant pulling velocity of 250 nm/s.

### Force-extension curve fitting and analysis

Force-extension curves describe the behavior of the tether during the stretch-and-relax cycles and were analyzed using a hybrid worm-like chain (WLC) polymer elasticity model based on Marko-Siggia’s WLC model (*13*) with modifications to incorporate enthalpic stretching (*47*). When a synaptic complex is formed the force is predominantly applied to dsDNA and the very stiff synaptic complex, therefore we initially model the force-extension behavior using the extensible WLC model (eq. 1) (*27, 47*):

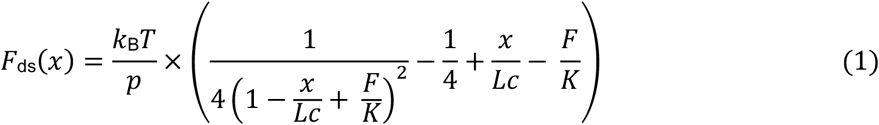

where *p* is persistence length, *K* is stretch modulus, and *Lc* is the contour length of the dsDNA and *x* – its extension. *k*_B_ is Boltzmann constant and *T* is room temperature. After synaptic complex disassembly, the construct contains additionally a flexible 114 nt long ssDNA, which requires the use of a hybrid WLC (hWLC) chain model describing the dsDNA and ssDNA flexibility with different WLC parameters (*13*). The ssDNA WLC term (eq. 2) uses *p*_ss_ = 2 nm (*13*), *L*_ss_ as the contour length of the ssDNA, and *x*_ss_ represents extension.

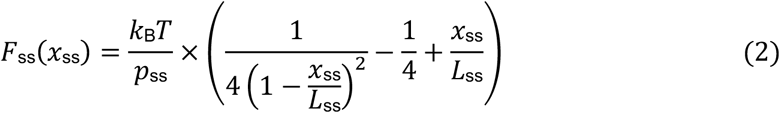

The total construct extension was calculated after WLC inversion using Cardano’s approximation

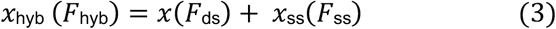

and the data was fitted using the hybrid force-extension behavior. Here, the contour length change from the initial dsDNA handle to the hybrid WLC construct was used as a fingerprint to verify complete synaptic complex disassembly (Fig. S3) based on the expected molecular fingerprint. Similarly, we identified subsequent combined hairpin unbinding and unfolding events showing the expected contour length increase up to 82 or 174 nm, depending on the exact *attC* sequence.

### Disassembly and unfolding force analysis

Data analysis was performed using Python 3.10.5 in Jupyter Notebook using common Python packages (Table S5) and lumicks.pylake to extract and analyze data from the C-trap. Briefly, high-frequency force, as well as piezo distance data (78 kHz), was downsampled by a factor of 300. We plotted force-extension curves (FECs) using the force signal of the immobile Trap 2 versus the extension (Piezo Distance) data. FECs were corrected for trap crosstalk. The measurements consisted of pulling and relaxing cycles, always reaching at least 30 pN force to ensure full hairpin unfolding. Using the relaxation trace as the most unfolded state in the presence of the protein the pulling traces were classified according to the observed integrase events using hWLC fitting and molecular fingerprints.

Synaptic complex disassembly events were identified as a peak on the FEC that led to changing states from synaptic complex to either integrase-bound hairpins or some of hairpin unfolding conformations. Similarly, hairpin unbinding forces were recorded for cases when the tether demonstrated no previous synaptic complex formation. We applied a minimum threshold of 6 pN to distinguish between free hairpin unfolding (5.7 pN, (*13*)) and integrase unbinding from the hairpins. All plots were generated using matplotlib or seaborn packages.

To compare the means of multiple datasets we used ANOVA (analysis of variance) that compares variance between and within the datasets. The analysis was done using Python packages such as pandas, pingouin, and statsmodels. With significantly different (P-value < 0.05) results the additional pairwise T-test with Holm correction was performed (allows to control the total error rate across all performed tests (*32*)). The pairwise T-test was used to identify an outlier dataset that differed from the group or to perceive the difference between two specific datasets.

### Protein mutagenesis

Protein variants were constructed using a QuikChange site-directed mutagenesis kit (Agilent, USA). The primer that introduced desired nucleotide changes was designed using QuikChange online tool (Table S3C) and the reaction was carried out as recommended by the manufacturer using amplification and subsequent digestion of template plasmid DNA with DpnI (Agilent, USA). A pMAL-c5X based plasmid that contained IntI1 fused with the maltose binding protein (MBP) was used as a template (*25*). Point mutations were confirmed using Sanger sequencing.

### Protein expression and purification

Plasmids containing mutated proteins were transformed into the *E. coli* BL21 (DE3) strain using heat-shock method. Using 5 mL of the overnight culture 1 L of Terrific broth (TB) medium with 50 μg/mL Ampicillin was inoculated to grow at 37°C and 140 rpm. Upon optical density = 0.6, protein expression was induced by adding Isopropyl-D-1-thiogalactopyranoside (IPTG) to a final concentration of 0.3 mM. The induced culture was grown at 14°C overnight and then centrifuged (Beckmann Coulter, USA) at 5500 rpm for 30 min at 4°C. The pellet was re-suspended, and cells were lysed using EmulsiFlex-C3 (Avestin, USA) and centrifuged at 8400 rpm for 1 hour at 4°C. The recovered lysate (supernatant) was filtered and loaded on the fast protein liquid chromatography (FPLC) device (ÄKTA Protein Purification System, Cytiva Life Sciences, USA). First purification step was using an amylose column (MBPTrap HP, 1 x 1 mL, Cytiva Life Sciences, USA) that was equilibrated beforehand according to the manufacturer’s instructions. Our MBP-tagged mutant protein was eluted using gradient in a maltose-containing buffer. The resulting fractions were checked using the sodium dodecyl sulfate polyacrylamide gel electrophoresis (SDS-PAGE) with 12% acrylamide concentration. In a second purification step, the remaining impurities were removed using ion exchange chromatography (HiTrap SP HP, 1 mL, Cytiva Life Sciences, USA). The resulting fractions were confirmed using 12% SDS-PAGE, dialyzed into storage buffer, snap-frozen, and stored at -70ºC (see *Supplementary Protocols*).

### Non-radiolabeled electrophoretic mobility shift assay (EMSA) and analysis

We developed an EMSA protocol, which allowed us to determine integrase binding to double- *attC* DNA constructs. To stabilize both *attC* hairpins and prevent Integrase binding to the long ssDNA spacer, we annealed two complementary staple oligos (Table S3F) to the double-*attC* construct (Fig. S5). 2 pmol of the DNA probe was added to the binding reaction in 15 μL volume with integrase (0-640 nM) and binding buffer (10 mM Tris-HCl, 10 mM NaCl, 40 mM KCl, 1 mM MgCl_2_, 4 mM EDTA, 1 mM DTT, 0.5 mg/mL BSA, 5% glycerol). Samples were incubated at room temperature for 15 min, then 2 μL of OrangeG dye was added and the samples were loaded to a 5% native polyacrylamide gel (acrylamide/bisacrylamide 37.5:1) containing 2.5% glycerol and 0.5x TBE (pre-run at 80 V for 1 hour, 4°C). The gel was run in 0.5x TBE, 2.5% glycerol buffer for 2 hours at 90 V at 4°C and subsequently stained with SYBR™ Gold (Thermo Fischer Scientific, USA) for 30 min in the dark on an orbital shaker. The gel was imaged using Azure300 Imager (Azure Biosystems, USA) in an epi-blue regime (460-490 nm wavelength) to visualize both free DNA and shifted DNA-protein complexes.

Image analysis was performed using Fiji (*48*) and its integrated gel analysis tool providing intensity values for bound states (shifted bands) and remaining free DNA for different protein concentrations. The fraction of bound species *b* was identified (eq. 4) and plotted against IntI1 concentration.

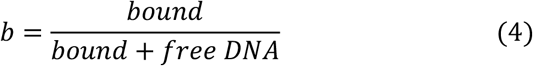

The data was fitted with the Hill-Langmuir equation (eq. 5) using MATLAB and the *C*_1/2_ and *b*_*max*_ (maximum bound fraction).

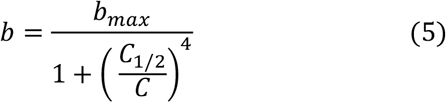

### The in vivo deletion assay using conjugation

This *in vivo* deletion assay is based on the suicide conjugation assay described by Bouvier et al. (*10*). This assay mimics the natural conditions in which cassettes carried by mobile integrons are delivered through horizontal gene transfer. Synthetic cassettes have been cloned in a pSW suicide vector (Table S3H-I). From the donor *E. coli* ß2163 strain, the pSW plasmid is delivered in a single-stranded form into a recipient *E. coli* MG1655 Δ*recA* strain. The pSW plasmid contains an RP4 origin of transfer (*oriT*RP4) oriented in such a way as to deliver either the reactive bottom strands of the *attC* recombination sites or the top ones. The synthetic cassette is inserted between a P_*tac*_ promoter and a promoter-less *pir116*\* gene that encodes a functional Π protein ([P_*tac*_]-*attC*-*lacI*^*q*^-*attC*-*pir116*\*). In the native [P_*tac*_]-*attC*-*lacI*^*q*^-*attC*-*pir116*\* configuration, the Π protein cannot be expressed and the pSW vector cannot be maintained in the recipient strain. However, if the cassette is deleted through an *attC* × *attC* recombination event catalyzed by IntI1, the *pir116*\* gene becomes expressed from the P_*tac*_ promoter, and the produced Π protein is able to sustain pSW replication in the recipient strain, which can be selected based on the pSW Cm^R^ marker. Note that *recA*-deficient recipient strain was used to limit the homologous recombination process between two identical *attC* sites during the conjugation step transfer.

Briefly, the donor strains were grown overnight in LB media supplemented with chloramphenicol (Cm) (resistance of the pSW plasmid), kanamycin (Km) (resistance of the ß2163 strain) and DAP (since ß2163 donor strain requires DAP to grow in rich medium), the recipient strain was grown overnight in LB media supplemented with spectinomycin (Sp) (resistance of the integrase expressing plasmid) and glucose (Glc, to repress the integrase gene when pBAD promoter is used). Both donor and recipient overnight cultures were diluted 1/100 in LB with DAP or arabinose (Ara) respectively and incubated until OD=0.7-0.8. 1 mL of each culture was then mixed and centrifuged at 3500 g for 6 minutes. The pellet was suspended in 100 μL LB, spread on a conjugation membrane (mixed cellulose ester membrane from Millipore, 47 mm diameter, and 0.45 μm pore size) over an LB+agarose+DAP+Ara Petri dish and incubated 3 hours for conjugation and recombination to take place. The membrane with the cells was then resuspended in 5 mL LB, after which serial 1:10 dilutions were made and plate on LB+agarose media supplemented with appropriate antibiotics. The recombination frequency was calculated as the ratio of recombinant CFUs, obtained on plates containing Cm, Sp and Glc, to the total number of recipient CFUs, obtained on plates containing only antibiotics corresponding to recipient cells (Sp and Glc). *attC* × *attC* recombination was confirmed by PCR using the “Sw23begin” and “Sw23end” primers (Table S3D). The recombination point was precisely determined by sequencing using the same primers. When we did not detect any recombinant CFUs, we considered that we obtained only one recombinant CFUs and we calculated that we called, the limit of detection (-). It corresponds to 1 recombinant CFUs / the total number of recipient CFUs obtained for all the n replicates. Note that in this case, we cannot calculate the standard deviation, so the bar chart does not show an error bar and the calculation of the *p-value* is not applicable (na).

It should be noted that in experiments using the same *attC* sites, we detected a few recombination events without integrase, while in others we detected none. This is due to recombination by microhomologies. This type of homologous recombination is independent of *recA* (*34*) and probably enhanced by the delivery of donor plasmids in single-stranded form (*i*.*e*., by conjugation). Therefore, a systematic integrase-free control was performed to establish the presence and frequency of such microhomology-driven recombination events (*33*).

## Acknowledgements

We thank all members of the Schlierf lab for lively discussions during the development of this project and in particular César Augusto Quintana-Cantaño for inspiring discussions and Dr. Yujun Zhang for supporting protein mutagenesis and production. This research was funded by TU Dresden core funds (M.S.), the Graduate Academy of TU Dresden (E.V.), supported by the Institut Pasteur, the Centre National de la Recherche Scientifique (CNRS-UMR 3525), ANR Chromintevol (ANR-21-CE12-0002-01), and by the French Government’s Investissement d’Avenir program Laboratoire d’Excellence ‘Integrative Biology of Emerging Infectious Diseases’ (ANR-10-LABX-62-IBEID) (D.M.). We acknowledge support by Dr. Jens Ehrig of the Tailored Smart Microscopy facility, a core facility of the EXC PoL at TU Dresden.

## Author Contributions

M.S. and E.V. conceptualized the study, M.S., C.L., D.M., and E.V. devised and validated the methodology, E.V. and C.L. performed the investigation, M.S. and D.M. supervised the project, E.V. and M.S. wrote the original draft and all other authors reviewed and edited it. M.S., E.V. and D.M. acquired funding for the project.

