## Supplementary Information for "The recombination efficiency of the bacterial integron depends on the mechanical stability of the synaptic complex"

### Supplementary Text

#### High affinity of IntI1 to non-specific ssDNA

Integrase binding behavior on different DNA constructs is largely unknown. There were many studies regarding specifically recognized DNA substrates, such as *attI* sites or *attC* sites as well as how the sequence-structure interplay affects integrase binding (*1–5*). However, what affinity does integrase have to the unspecific sequence, does it recognize nucleotide motifs of the *attI* or does it unspecifically bind the double-stranded DNA and perform diffusion/sliding on the molecule to bind the *attI* is unknown. We aimed to gain some initial understanding of the mode of integrase recognition and binding to an unspecific DNA substrate. We used a correlative optical tweezers setup (C-trap, LUMICKS) that has a confocal microscope to be used simultaneously with optical trapping. A DNA molecule, functionalized at the ends with biotins was tethered to the streptavidin-coated polystyrene beads and stretched to introduce small tension as well as to expose the sequence to the solution. We used our engineered fusion integrase protein that carries a monomeric enhanced green fluorescent protein ($IntI1_{mEGFP}$) to investigate how IntI1 binds the unspecific DNA substrate.

We first used a full λ-phage DNA sequence as the substrate. This DNA molecule is long (48.5 kbp) allowing for better imaging conditions and has several motifs that mimic the *attI* site. The *attI* binding box sequence allows variability, the consensus is six to seven nucleotides with conserved GTT triplet followed by three to four purines and one pyrimidine base (*6*). Such sequence flexibility results in a higher probability of finding such binding motifs on a long DNA molecule. In fact, on the λ-phage DNA, the GTTRRRRY motif is found 29 times. We tethered the molecule (biotinylated double-stranded DNA, LUMICKS) to the beads and stretched to 20 pN to expose the sequence to the $IntI1_{mEGFP}$ protein (C = 50 nM). However, no binding was detected. We repeated the same procedure on another DNA construct of a shorter length (20452 bp, biotinylated single-stranded DNA, LUMICKS) with biotin functionalization on the ends of the same DNA strand. This type of attachment provides the possibility to transform the dsDNA into a ssDNA while already in the sample chamber of the setup. The DNA was attached again to the streptavidin-coated beads and then overstretched to melt away one of the strands. Once melted, the molecule that remains tethered to the beads is purely single-stranded. We exposed such molecule to 50 nM $IntI1_{mEGFP}$ first in its dsDNA state – and again saw no binding. Then we overstretched the molecule to produce a ssDNA and exposed it again to the same $IntI1_{mEGFP}$ solution. The results were unexpected, as we observed multiple binging events along the whole length of the ssDNA molecule (Fig. S9).

The binding occurred in a wide range of forces, from 5 pN to 40 pN, and was quite durable (over 2.5 minutes). A bound $IntI1_{mEGFP}$ monomer has remained bound even upon the stretching of the molecule in the indicated range. We observed no detectable translocation of the protein along the DNA molecule based on recorded kymographs. We used the same DNA construct to probe how integrase would behave in the presence of the hybrid, overstretched DNA. We first exposed the dsDNA molecule to the $IntI1_{mEGFP}$ solution with a small tension of 7 pN, again we observed no integrase binding. Then we started to overstretch the tether with small stalls at 15 and 30 pN to allow for some interaction, however, none was observed. We finally brought the tension to 65 pN and saw the beginning of DNA melting with occasional release of tension due to the appearance of local openings of the DNA strands. After a minute of the melting process, keeping the tether at ~65 pN we observed first $IntI1_{mEGFP}$ binding events. Those appeared on the tether closer to the beads, corresponding to the opening of melted DNA bubbles along the molecule. Some interactions were again very stable, $IntI1_{mEGFP}$ remained bound at 65pN DNA tension for over two minutes (Fig. S9B). Once the tether broke, we had no more binding signal from the $IntI1_{mEGFP}$ between the beads. With this preliminary data on the integrase binding behavior, we discovered that it has a surprisingly high affinity to unspecific ssDNA, independent of the tension forces acting at the molecule. On another hand, we observed, unexpectedly, no binding of $IntI1_{mEGFP}$ to the unspecific as well as *attI*-like dsDNA sequences.

***Mechanical work during synaptic complex disassembly***

We have characterized the stability of synaptic complex using disassembly force and found that lower efficiency *attC* sites showed at least 5 pN lower mean disassembly force. This is approximately 0.6 times lower than the reference high-efficiency $attC^{bs}_{aadA7}$. To better present the difference between the biochemical and biophysical processes that occur in the cell we aimed to assess the energetics of the transition from a synaptic complex to its disassembled state (the hairpin-bound state). We calculated the total mechanical work ($\Delta W = \Delta W_{dissipated} + \Delta W_{reversible}$) for all investigated protein-DNA combinations and presented the results in biologically relevant units of $k_BT$. The mechanical work provides us with insights on the energy difference between two stable states in our non-equilibrium experiments; the synaptic complex and the hairpin-bound state. As we assume the latter to have the same Gibbs free energy for all our *attC* sites, $\Delta W$ manifests into the different stabilities of the synaptic complex. With more work during the transit from the synaptic assembly to the hairpin-bound state, we infer a more energetically favorable synaptic complex assembly (a state with lower Gibbs free energy). Thus, using mean mechanical work ($\overline{\Delta W}$) we have built an energy profile with varying stabilities for synapses containing different *attC* sites or mutant variants.

All $\overline{\Delta W}$ values remain in a range of 62-124 $k_BT$, which suggests that there is a large amount of dissipation ($\Delta W_{dissipated}$) This is a hallmark for hysteresis which has several implications in biology out of equilibrium. As expected from the force difference, the low recombination efficiency *attC* sites did not vary much between themselves in terms of $\overline{\Delta W}$ ranging from 78 $k_B$T to 64 $k_B$T for $attC^{ts}_{aadA7}$ and $attC_{VCR2/1}$ (Fig. S10C). Interestingly, a synaptic complex containing a highly efficient $attC^{bs}_{aadA7}$ and IntI1 required 1.5 times more energy than its counterpart $attC^{ts}_{aadA7}$. However, exchanging the protein to the C-terminal deficient IntI1$^{\Delta C}$ decreased the work of the disassembly twice ($\overline{\Delta W}$ = 62 $k_B$T), likely increasing the Gibbs free energy of the synaptic formation and making it less favorable. Very close work values were obtained for low recombinogenic $attC_{VCRinv}$ ($\overline{\Delta W}$ = 67 $k_B$T), that differs four orders of magnitude from the $attC^{bs}_{aadA7}$ *in vivo*. We must note that as the C-terminal deletion can theoretically influence the hairpin-bound formation by destabilizing the dimer assembly on the *attC*, the above comparison should be taken with caution. We cannot be certain that the hairpin-bound state has the same Gibbs free energy for the integrase mutant due to altered protein interactions. Alanine mutant IntI1$^{A}$ has mutations that are not expected to manifest in the hairpin-bound state as the αI2 helix residues that were changed are involved in the *attC* interaction *in trans* and do not play a role in the hairpin binding alone.

The mechanical work (*W*) was calculated as the integral of force (*F*) function over displacement (*x*) (eq. 1).

$$W = \int_{x0}^{x1} F(x)dx \qquad (1)$$

For this, an hWLC was plotted for a synaptic complex-bound state (S) and the next stable hairpin-bound state (HB). The integral was calculated for both curves separately for *x* in the range from 0 to the distance of the disassembly event ($x_1$) producing $W_S$ and $W_{HB}$. The work done by the tether upon synaptic complex disassembly was calculated by subtraction of the work done by the tether from the HB state forth ($W_{HB}$) from the synaptic complex state work $W_S$. This $\Delta W$ is the synaptic complex disassembly work (Fig. S10) that depends on the $F_{diss}$ (synaptic complex disassembly force). Every measuring condition (the combination of an *attC* and a protein variant) has its characteristic unfolding force dataset that we use to obtain the corresponding characteristic work values. It was recalculated to $k_B$T to make it more relevant in the context of biological processes. Free energy profiles were constructed as a model representation using synaptic complex disassembly work ($\Delta W$) as an energy term of transition between the states (*7*). The reaction coordinate is arbitrary.

### Supplementary Figures:

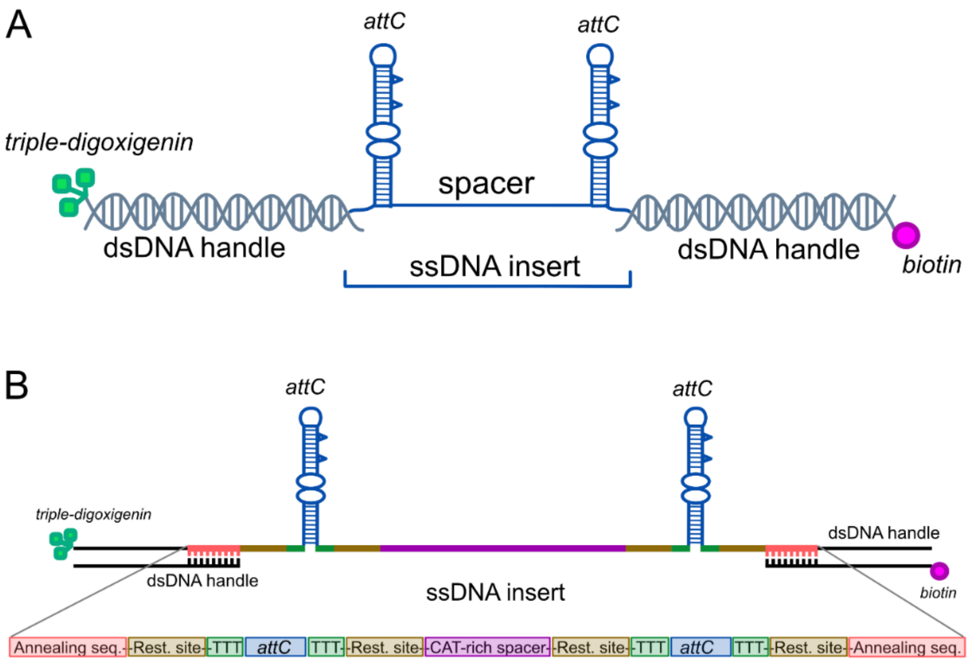

**Figure S1. Scheme of the hybrid DNA tether assembly and composition. (A)** Hybrid DNA construct for *in vitro* synapse reconstitution: two *attC* hairpins, a spacer for gene cassette imitation, dsDNA handles with 5' modifications carrying biotin and triple-digoxigenin for specific bead attachment in the optical tweezers assay. **(B)**. Schematic representation of the single-stranded insert. Two *attC* hairpins are indicated as well as the ssDNA spacer. The position of three thymine bases (TTT), restriction sites (Rest. site), the CAT-rich spacer, and complementary overhangs (Annealing seq.) are labeled and indicated with colored rectangles.

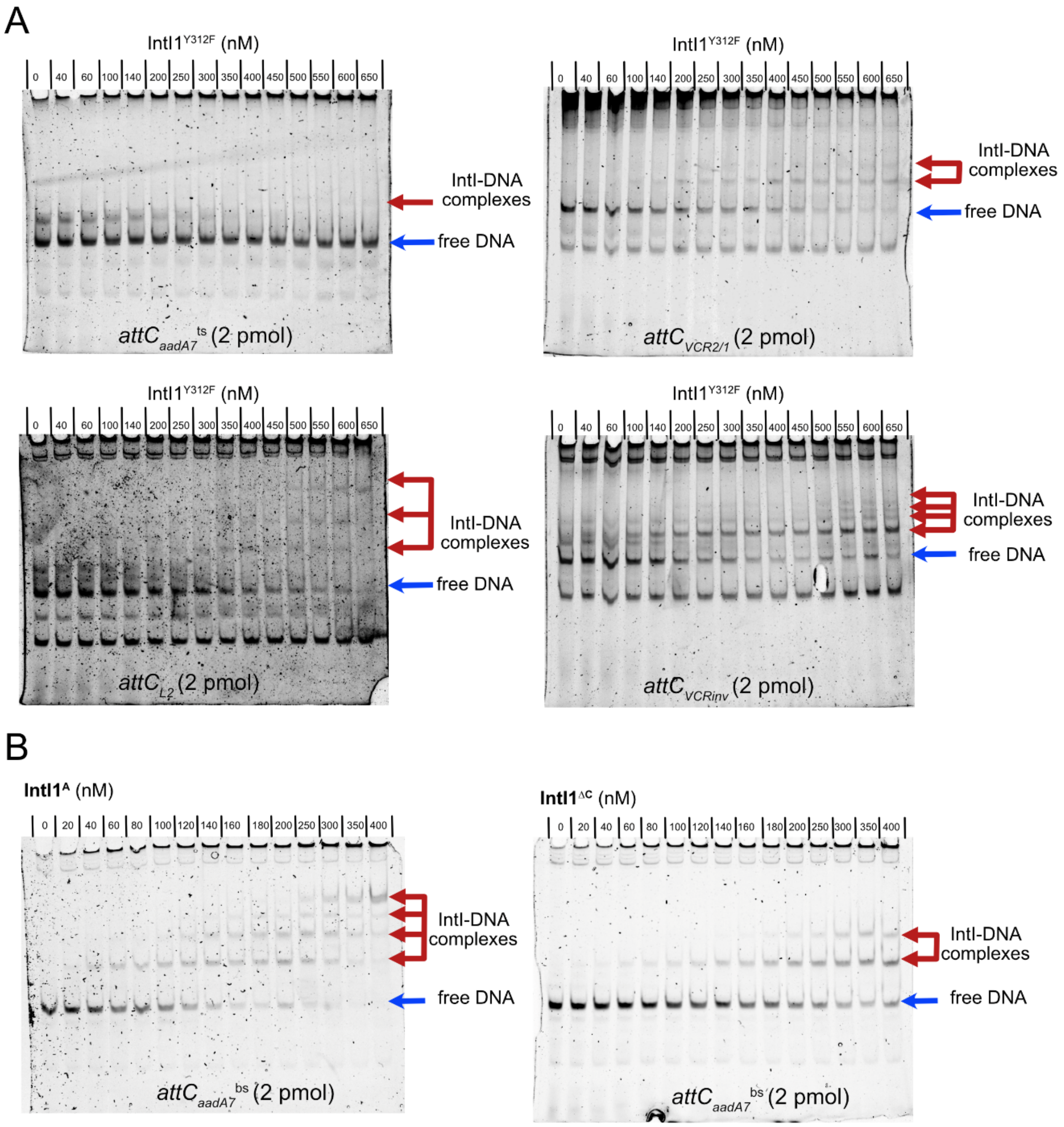

**Figure S2. EMSA of different *attC* sites with IntI1 variants. (A)** EMSA with IntI1 on $attC^{ts}_{aadA7}$, $attC_{L2}$, $attC_{VCR2/1}$, and $attC_{VCRinv}$ showed from one to four integrase subunit binding modes. **(B)** EMSA with IntI1$^{A}$ and IntI1$^{\Delta C}$ on $attC^{bs}_{aadA7}$ showed from one to four subunit binding modes.

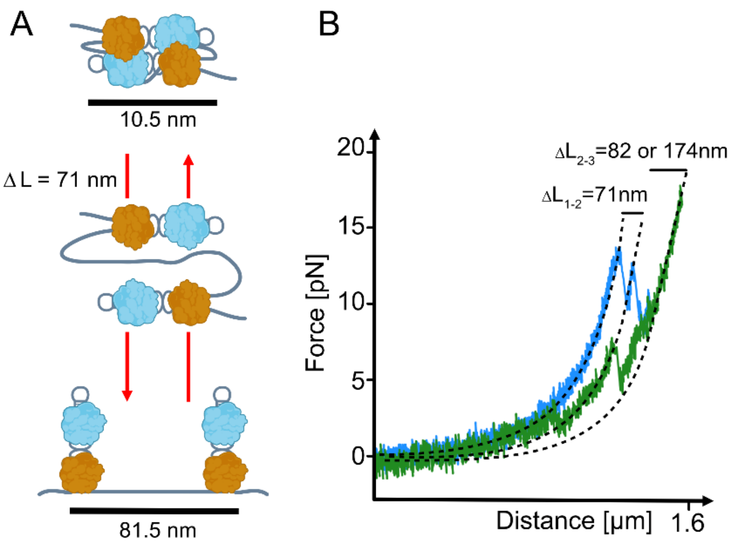

**Figure S3. Scheme of the synaptic complex disassembly molecular fingerprint on the double-*attC* tether. (A)** Synaptic complex disassembly scheme with the characteristic contour length change of 71 nm. **(B)** Characteristic force-extension curves for high (in blue) and low (in green) recombination efficiency *attC* sites. Synaptic complex disassembly is identified using an established molecular fingerprint $\Delta L_{1\text{-}2}$=71 nm (experimental data $\Delta L_{1\text{-}2}$=71 ± 2 nm, N=38), and the characteristic force of the event is recorded using the breaking point of the state (1) on the force-extension curve. Subsequent events of hairpins unfolding produce the theoretical $\Delta L_{2\text{-}3}$=82 nm for short *attC*s or 174 nm for VCR *attC* sites (experimental data $\Delta L_{2\text{-}3}$= 83 ± 2 nm, N=11 and 174 ± 1 nm, N=5).

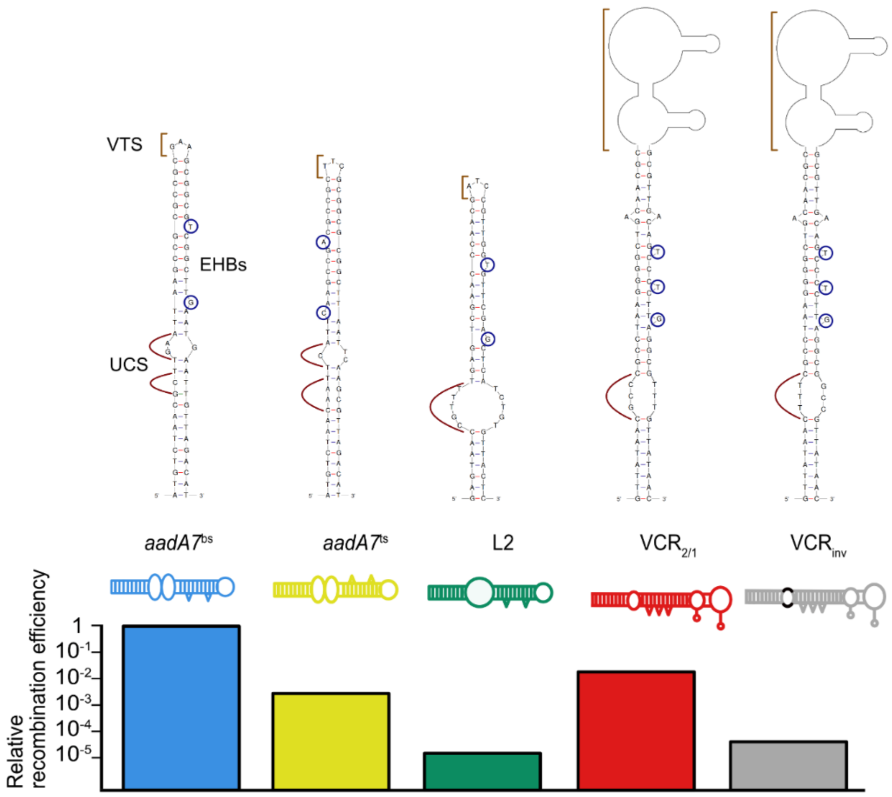

**Figure S4. Sequence and folding architecture of the investigated *attC* sites depicted using Mfold** (*8*). Structural parts are indicated: UCS with red lines, EHBs with blue circles, and VTS with brown lines. Relative recombination efficiency of the integration *in vivo* (*2*, *4*, *5*, *9*, *10*).

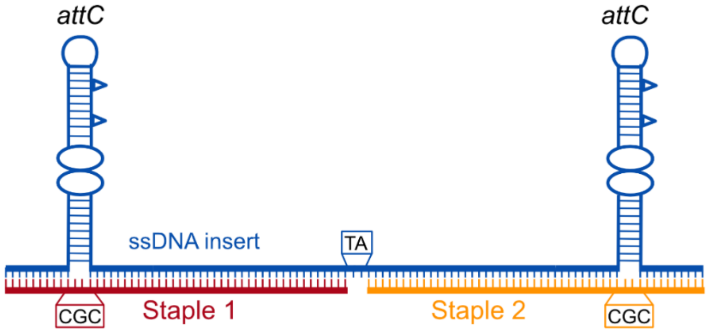

**Figure S5. EMSA sample preparation design to minimize the amount of ssDNA.** We stabilized the hairpin structure using two long complementary oligonucleotides (staple 1 and staple 2). CGC sequence was inserted to accommodate the width of the hairpin, TA bases in the middle of the spacer were left unpaired to allow the spacer necessary flexibility for synaptic complex assembly.

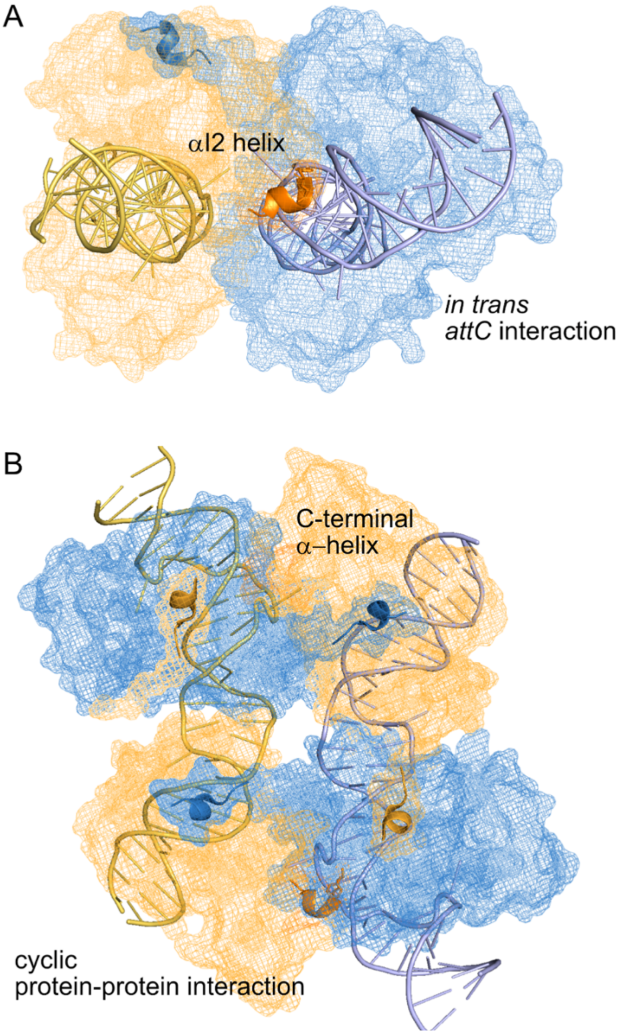

**Figure S6. Protein-DNA and protein-protein interactions that are specific only for the synaptic complex formation. (A)** αI2 helix interaction with adjacent DNA backbone *in trans* by $K^{209}$ and $Y^{210}$ (numbering according to IntI4 resolved synapse, *(11)*). **(B)** C-terminal α-helices that cyclically connect subunits in a tetramer. Visualized with PyMOL *(12)*.

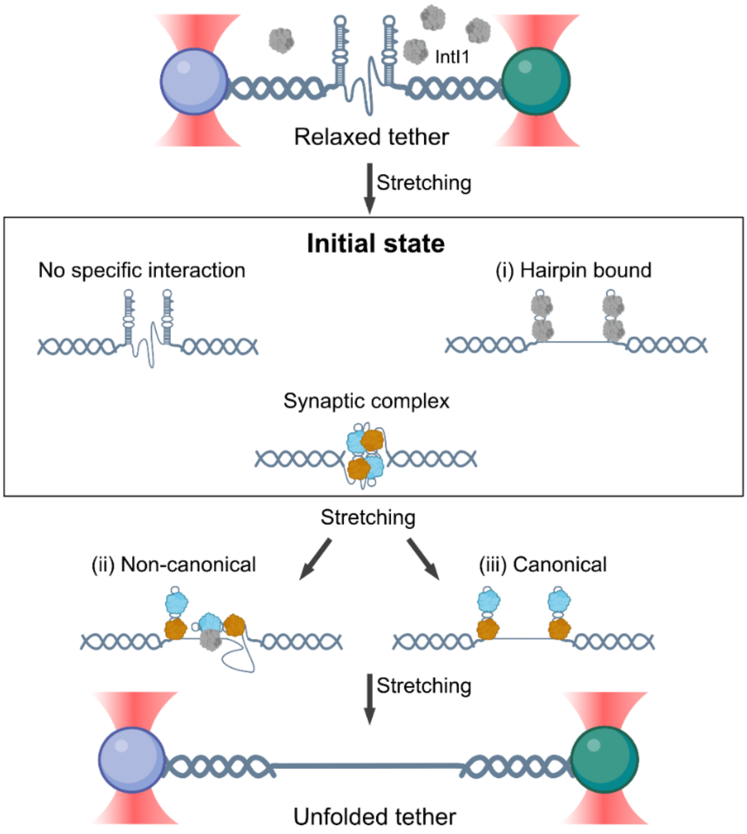

**Figure S7. Schematic representation of possible integrase interaction modes with the double-*attC* tether.** Three possible initial states resulting from integrase interaction with the tether are depicted in a frame: no specific interaction, (i) hairpin-bound or synaptic complex. Synaptic complex initial state can further disassemble via two pathways leading to (ii) non-canonical or (iii) canonical structures.

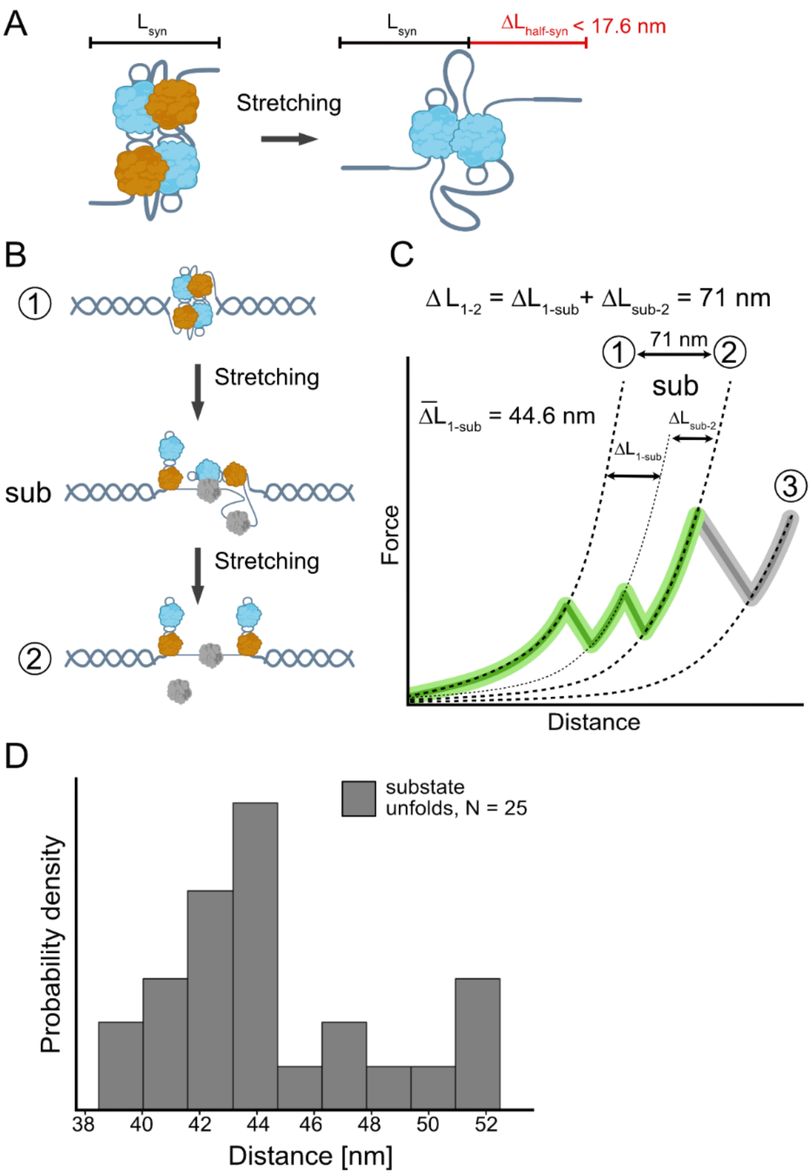

**Figure S8. A scheme of possible non-canonical substate structural formations. (A)** A schematic representation of the maximum contour length change possible for a semi-formed synaptic complex (half-syn). The ΔL < 17.6 nm is the maximum value achieved by unfolding both R-boxes (26 nts). **(B)** A schematic representation of the proposed non-canonical pathway substate structure formation upon the stretching of the tether. **(C)** A force-extension curve depicting the substate and the mean contour length change to the substate ($\Delta L_{1\text{-}sub}$) and from the substate to the hairpin-bound state ($\Delta L_{sub\text{-}2}$). **(D)** A histogram of the contour length change for non-canonical unfolding events that led to the substate structure ($\Delta L_{1\text{-}sub}$), sampled from different *attC* sites.

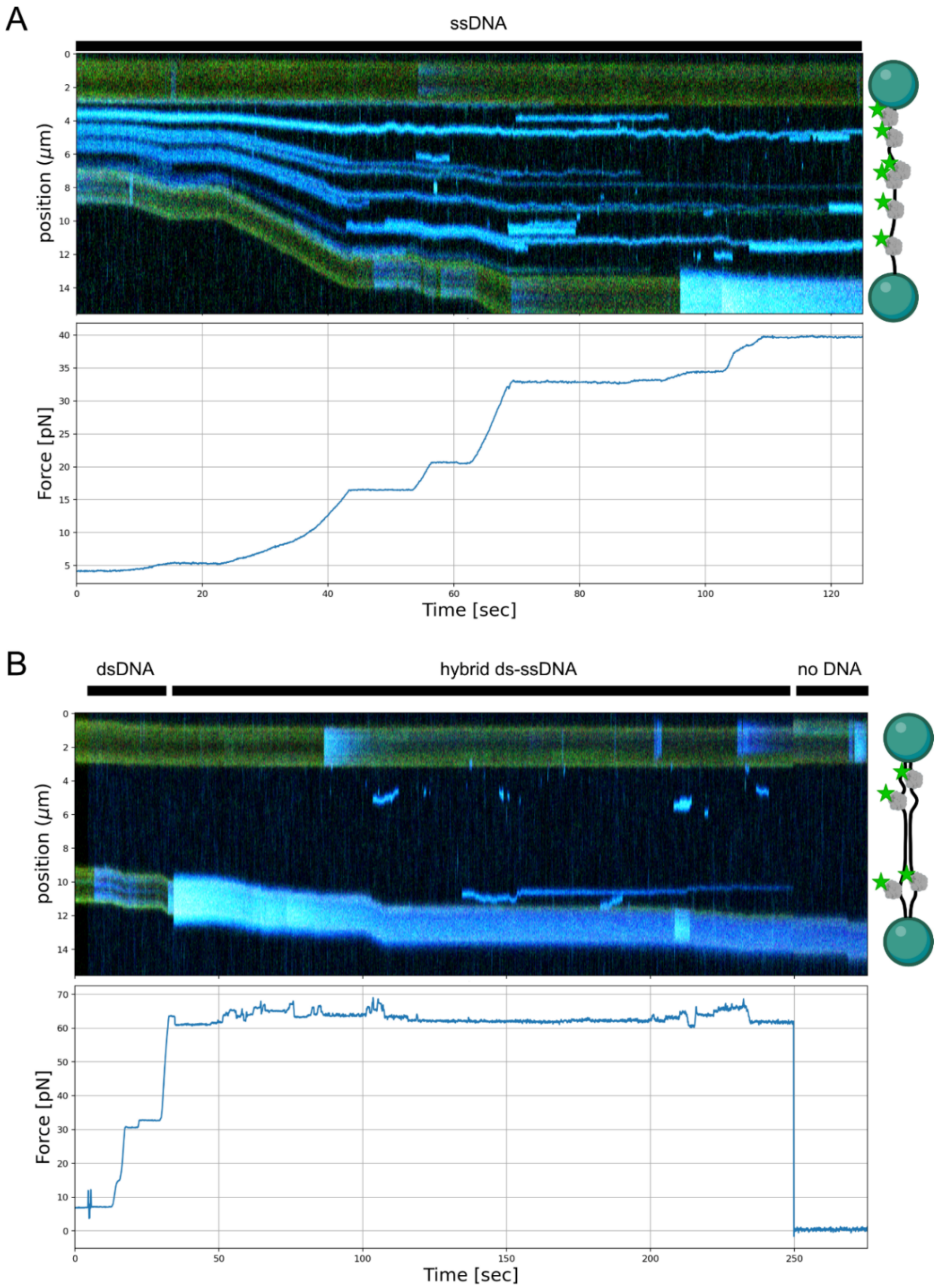

**Figure S9. Kymographs and corresponding forces of the ds-ssDNA tethers in the presence of IntI1$_{\mathbf{mEGFP}}$. (A)** ssDNA tether is stretched from 5 to 40 pN in the presence of IntI1$_{mEGFP}$. **(B)** The dsDNA molecule is overstretched to a hybrid ds-ssDNA state in the presence of IntI1$_{mEGFP}$. The blue signal indicates integrase binding.

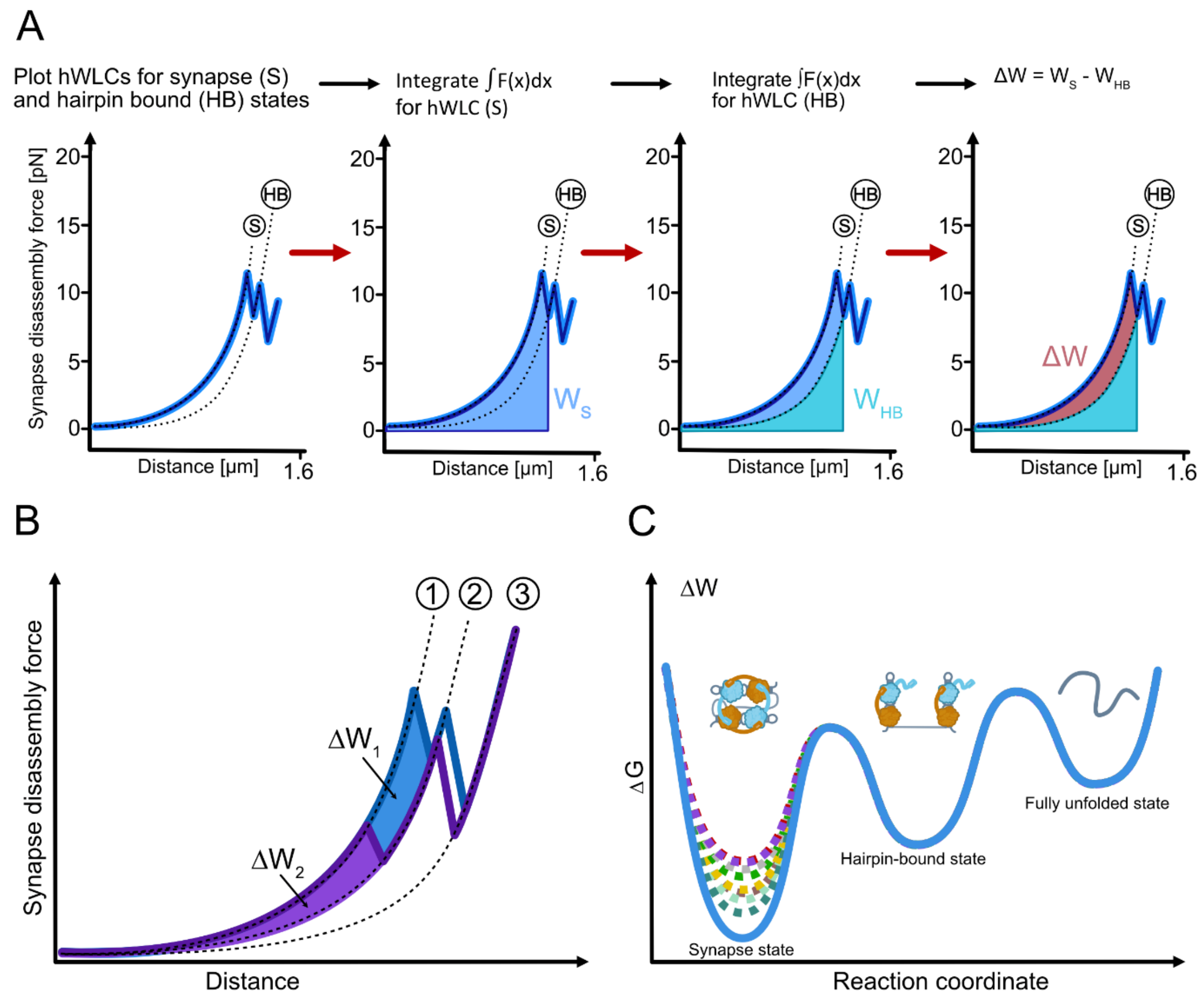

**Figure S10. Mechanical work calculations and free energy profile as a model representation of varying synaptic complex stability between different *attC* sites and protein variants. (A)** Schematic representation of synaptic complex mechanical work calculation (ΔW). First, we plotted the hWLC of the synapse and hairpin-bound states, then we integrated the area under the synapse state curve – $W_S$. Next, we integrated the area under the hairpin-bound state curve – $W_{HB}$ and subtracted it from the $W_S$ to get only the mechanical work done by the tether upon synaptic complex disassembly – ΔW. **(B)** Mechanical work of synaptic complex disassembly for high ($\Delta W_1$) and low ($\Delta W_2$) mechanical stability. **(C)** Free energy profile of transition between synaptic complex, hairpin-bound and fully unfolded state determined from the mean mechanical work upon synapse disassembly ($\overline{\Delta W}$) for $attC^{bs}_{aadA7}$ (blue), $attC^{ts}_{aadA7}$ (yellow), $attC_{L2}$ (green), $attC_{VCRinv}$ (grey), $attC^{bs}_{aadA7}$ × $attC_{VCR2/1}$ (brown), $attC^{bs}_{aadA7}$ × $attC_{L2}$ (mint), $IntI1^{\Delta C}$ (purple), $IntI1^{A}$ (teal).

### Supplementary Tables

**Table S1. Relative recombination efficiencies *in vivo* and synapse disassembly force for investigated IntI1:double-*attC* combinations.** Relative recombination efficiency of the integration events for $attC_{aadA7}$ (both strands) is taken from (*9*, *10*), for $attC_{L2}$ from (*4*), for $attC_{VCR2/1}$ from (*9*), for $attC_{VCR\mathrm{inv}}$ from (*2*, *5*); nr – no IntI1-mediated recombination. Calculated median, mean force, and the SEM (standard error of the mean) are presented for all synaptic complex disassembly events.

| *attC* / IntI1 variant | *aadA7* $^{bs}$ | *aadA7* $^{ts}$ | $VCR_{2/1}$ | *aadA7* $^{bs}$ x $VCR_{2/1}$ | L2 | $VCR_{inv}$ | *aadA7* $^{bs}$ x L2 |
|---|---|---|---|---|---|---|---|
| | IntI1 | | | | | | |
| Relative recombination efficiency (integration) | 1 | $5.38*10^{-3}$ | $3.55*10^{-2}$ | - | $2.5*10^{-5}$ | $7.56*10^{-5}$ | - |
| Relative recombination efficiency (excision) | 1 | $1.03*10^{-2}$ | - | $4.96*10^{-2}$ | $3.55*10^{-4}$ | $3.69*10^{-4}$ | nr |
| Median force (pN) | 12.7 | 8.2 | 6.3 | 7.9 | 7.5 | 7.1 | 9.4 |
| Mean force (pN) | 13.9 | 8.8 | 7.1 | 9.0 | 8.0 | 7.6 | 11.4 |
| SEM (pN) | 0.6 | 0.4 | 0.3 | 0.4 | 0.3 | 0.4 | 0.6 |

| *attC* / IntI1 variant | *aadA7* $^{bs}$ | |
|---|---|---|
| | $IntI1^{A}$ | $IntI1^{\Delta C}$ |
| Relative recombination efficiency (integration) | - | - |
| Relative recombination efficiency (excision) | nr | nr |
| Median force (pN) | 9.4 | 5.6 |
| Mean force (pN) | 11.4 | 6.8 |
| SEM (pN) | 0.6 | 0.4 |

**Table S2. Table of all measurement data.** The data collected in the integrase-filled channel is indicated as "IntI channel", non-specific integrase interaction events are indicated as well as the hairpin-bound events (HB) and synaptic complex disassembly events (ScD).

| **Protein** | ***attC*** | **# specific tethers** | **# traces with data (IntI channel)** |
|---|---|---|---|
| IntI1 | *aadA7* $^{bs}$ | 29 | 313 |
| | *aadA7* $^{ts}$ | 23 | 400 |
| | L2 | 22 | 314 |
| | $VCR_{2/1}$ | 22 | 251 |
| | *aadA7* $^{bs}$ x $VCR_{2/1}$ | 2 | 324 |
| | $VCR_{inv}$ | 19 | 155 |
| | *aadA7* $^{bs}$ x L2 | 4 | 306 |
| $IntI1^{A}$ | *aadA7* $^{bs}$ | 19 | 380 |
| $IntI1^{\Delta C}$ | *aadA7* $^{bs}$ | 25 | 385 |

| **Protein** | ***attC*** | **All channels** | **IntI channel** | | | | |
|---|---|---|---|---|---|---|---|
| | | **ScD** | **ScD** | | | **Non-specific** | **HB** |
| | | | **Total** | **Canonical** | **Non-canonical** | | |
| IntI1 | *aadA7* $^{bs}$ | 169 | 128 | 95 | 33 | 180 | 55 |
| | *aadA7* $^{ts}$ | 175 | 163 | 70 | 93 | 207 | 30 |
| | L2 | 151 | 146 | 57 | 89 | 156 | 12 |
| | $VCR_{2/1}$ | 126 | 124 | 50 | 74 | 104 | 22 |
| | *aadA7* $^{bs}$ x $VCR_{2/1}$ | 214 | 214 | 60 | 154 | 65 | 45 |
| | $VCR_{inv}$ | 75 | 74 | 12 | 62 | 71 | 10 |
| | *aadA7* $^{bs}$ x L2 | 200 | 200 | 65 | 135 | 106 | 50 |
| $IntI1^{A}$ | *aadA7* $^{bs}$ | 144 | 141 | 25 | 116 | 169 | 70 |
| $IntI1^{\Delta C}$ | *aadA7* $^{bs}$ | 139 | 129 | 44 | 85 | 195 | 61 |

**Table S3. DNA sequences used in this study. (A)** Primers for dsDNA handles. **(B)** Primers for dsDNA insert. **(C)** Primers for protein mutagenesis. **(D)** Primers and oligonucleotides for *in vivo* studies. **(E)** Full double-*attC* insert sequences. **(F)** Staple oligonucleotides for EMSA. **(G)** Plasmids. **(H)** Bacterial strains for *in vivo* conjugation studies. **(I)** Constructed plasmids for *in vivo* conjugation studies.

**A.** Primers designed for PCR amplification of dsDNA handles for double-*attC* constructs. Sequences are given in a 5' to 3' direction.

| **Biotin functionalized dsDNA handle from lambda-DNA template** | |
|---|---|
| biotin_fwd_852bp_handle_AM | [Biotin] - CAGCATTGGTGACCTTGTTC |
| bio_rev_2500bp_primer | GGTATAGCATTCTAGCATTCAGCAGCACTTTAAACTGTCG |
| **Triple-digoxigenin functionalized dsDNA handle from lambda-DNA template** | |
| Triple-dig_fwd (70) | [Triple-DIG] - ATCCGCAGAAGACGCAGATGCC |
| 3Dig_rev_2500bp_primer | ACGCAATGCGCAATGCAGGTCTTTTTCTGCTCTGACATGACG |

**B.** Primers designed for PCR amplification of dsDNA insert sequence using a corresponding plasmid template. 5'- phosphorylation is shown as [Phos]. Sequences are given in a 5' to 3' direction.

| **Insert with double-*attC*$_{aadA7}^{bs}$; double-*attC*$_{aadA7}^{ts}$** | |
|---|---|
| db_aadA7_few_sh | CATTGCGCATTGCGTAGATCTTTTATG |
| db_aadA7_rev_sh_phos | [Phos] - CTAGAATGCTATACCCTCGAGAAAATGTC |
| **Insert with double-*attC*$_{L2}$; double-*attC*$_{VCR2/1:}$ double-*attC*$_{VCRinv}$** | |
| Fwd_VCR | CATTGCGCATTGCGTAGATCTTTTG |
| Rev_VCR_phos | [Phos] - CTAGAATGCTATACCCTCGAGAAAG |

**C.** Primers for protein mutagenesis. Sequences are given in a 5' to 3' direction

| **Protein mutagenesis using QuikChange (Agilent)** | |
|---|---|
| primer_fwd_mut_stop | GTGCTGAAAGTTGGCGGTTAGTGAGTGCGCTCACCGCTTG |
| primer_fwd_mut_Ala | ACGCCCTTGAGCGGGCGGCTCCGCGCGCCGGGC |

**D.** Primers and oligonucleotides used for *in vivo* studies. Sequences are given in a 5' to 3' direction

| | |
|---|---|
| 7166 | CGTCGATGAAGATGAATTTTCTGGCG |
| 7167 | CGCCAGAAAATTCATCTTCATCGACG |
| 7168 | GCAATGGCATCCTGGTCATCCAGCGG |
| 7169 | CCGCTGGATGACCAGGATGCCATTGC |
| 7223 | TGAGGCAGCGCAAGTCAATCCTGGCGGATTCACTACCCCTGCGCGAAGGC |

| | |
|---|---|
| 7229 | AATTCGGTTATAACGGCCGCCTCAAGAGGGACTGTCAACGCGTGGCGTTTCCAGTCCCATTGAGCCGCGGTGG |
| 7230 | TTGCTGTTGTTGTGTTTGAGTTTAGTGGTAGTGCGTTGTCAGCCCCTTAGGCGAAAGTTAGATGG |
| 7231 | GAAACGCCACGCGTTGACAGTCCCTCTTGAGGCGGCCGTTATAACCG |
| 7232 | GATCCCATCTAACTTTCGCCTAAGGGGCTGACAACGCACTACCACTAAACTCAAACACAACAACAGCAACCACCGCGGCTCAATGGGACTG |
| 7233 | CGGATCGTTGGGTTCGACTCAAAACGGTTACTCGGTATTTGCATGCTTACAGGAGC |
| 7234 | TTGGTGTTCGAGCTCATCTGTGTTACTCGATGGTCTAGAGGCGAAGC |
| 7235 | AATTCGGTTAGAGTAACACAGATGAGCTCGAACACCAACGGATCGTTGGGTTCGACTCAAAACGGTTACTCGATGG |
| 7236 | GATCCCATCGAGTAACCGTTTTGAGTCGAACCCAACGATCCGTTGGTGTTCGAGCTCATCTGTGTTACTCTAACCG |
| 7237 | CGCGGCTCAATGGGACTGGAAACGCCACGCGTTGACAGTCCCTCTTGAGGCGGCCGTTAGATGGTCTAGAGGCGAAGCGGC |
| 7238 | GTGGTTGCTGTTGTTGTGTTTGAGTTTAGTGGTAGTGCGTTGTCAGCCCCTTAGGCGAAAGTTAAGGTATTTGCATGCTTACAGGAGC |
| 7245 | CTAACCGCCAACTTTCAGCACATGCGTGTAAATCATCGTCGTAGAGACGT CGG |
| 7246 | GCCCGCTCAAGGGCGTCGGGAAGCGC |
| 7247 | GGCTCCGCGCGCCGGGCATTCCTGGC |
| Swbeg | CCGTCACAGGTATTTATTCGGCG |
| Swend | CCTCACTAAAGGGAACAAAAGCTG |

**E.** Designed sequences of the insert in pUC57-Kan-Simple carrying vector amplified with the primers above. The sequence of an *attC* site is indicated in red; three dT are in grey; restriction sites flanking the hairpins are in green; ligation parts to the nicked handles are shown in blue and underlined. Sequences are given in a 5' to 3' direction.

| | |
|---|---|
| $attC_{aadA7}^{bs}$ | CATTGCGCATTGCGTAGATCTTTTATGTCTAACGCTTGAATTAAGCCGCGCCGCGAAGCGGCGTCGGCTTGAATGAATTGTTAGACATTTTCCATGGCTATATTTCAGCATCATCACATCATCATCATCATCATCACAGGGTAGATCATCATCATCAACACGGGACGACAGCAAATGGCACCCTTCTATAAGCTTTTTATGTCTAACGCTTGAATTAAGCCGCGCCGCGAAGCGGCGTCGGCTTGAATGAATTGTTAGACATTTTCTCGAGGGTATAGCATTCTAG |
| $attC_{aadA7}^{ts}$ | CATTGCGCATTGCGTAGATCTTTTATGTCTAACAATTCATTCAAGCCGACGCCGCTTCGCGGCGCGGCTTAATTCAAGCGTTAGACATTTTCCATGGCTATATTTCAGCATCATCACATCATCATCATCATCATCACAGGGTAGATCATCATCATCAACACGGGACGACAGCAAATGGCACCCTTCTATAAGCTTTTTATGTCTAACAATTCATTCAAGCCGACGCCGCTTCGCGGCGCGGCTTAATTCAAGCGTTAGACATTTTCTCGAGGGTATAGCATTCTAG |

| | |
|---|---|
| *attC*$_{L2}$ | CATTGCGCATTGCGTAGATCTTTTGAGTAACCGTTTTGAGTCGAACCCAACGATCCGTTGGTGTTCGAGCT<br>CATCTGTGTTACTCTTTCCATGGCTATATTTCAGCATCATCACATCATCATCATCATCATCACAGGGTAGA<br>TCATCATCATCAACACGGGACGACAGCAAATGGCACCCTTCTATAAGCTTTTTGAGTAACCGTTTTGAGTC<br>GAACCCAACGATCCGTTGGTGTTCGAGCTCATCTGTGTTACTCTTTCTCGAGGGTATAGCATTCTAG |
| *attC*$_{VCR2/1}$ | CATTGCGCATTGCGTAGATCTTTTGTTATAACGCCCGCCTAAGGGGCTGACAACGCACTACCACTAAACTC<br>AAACACAACAACAGCAACCACCGCGGCTCAATGGGACTGGAAACGCCACGCGTTGACAGTCCCTCTTGAGG<br>CGTTTGTTATAACTTTCCATGGCTATATTTCAGCATCATCACATCATCATCATCATCATCACAGGGTAGAT<br>CATCATCATCAACACGGGACGACAGCAAATGGCACCCTTCTATAAGCTTTTTGTTATAACGCCCGCCTAAG<br>GGGCTGACAACGCACTACCACTAAACTCAAACACAACAACAGCAACCACCGCGGCTCAATGGGACTGGAAA<br>CGCCACGCGTTGACAGTCCCTCTTGAGGCGTTTGTTATAACTTTCTCGAGGGTATAGCATTCTAG |
| *attC*$_{VCRinv}$ | CATTGCGCATTGCGTAGATCTTTTGTTATAACTTTCGCCTAAGGGGCTGACAACGCACTACCACTAAACTC<br>AAACACAACAACAGCAACCACCGCGGCTCAATGGGACTGGAAACGCCACGCGTTGACAGTCCCTCTTGAGG<br>CGGCCGTTATAACTTTCCATGGCTATATTTCAGCATCATCACATCATCATCATCATCATCACAGGGTAGAT<br>CATCATCATCAACACGGGACGACAGCAAATGGCACCCTTCTATAAGCTTTTTGTTATAACTTTCGCCTAAG<br>GGGCTGACAACGCACTACCACTAAACTCAAACACAACAACAGCAACCACCGCGGCTCAATGGGACTGGAAA<br>CGCCACGCGTTGACAGTCCCTCTTGAGGCGGCCGTTATAACTTTCTCGAGGGTATAGCATTCTAG |
| *attC*$_{aadA7}^{bs}$ x *attC*$_{VCR2/1}$ | CATTGCGCATTGCGTAGATCTTTTATGTCTAACGCTTGAATTAAGCCGCGCCGCGAAGCGGCGTCGGCTTG<br>AATGAATTGTTAGACATTTTCCATGGCTATATTTCAGCATCATCACATCATCATCATCATCATCACAGGGT<br>AGATCATCATCATCAACACGGGACGACAGCAAATGGCACCCTTCTATAAGCTTTTTGTTATAACGCCCGCC<br>TAAGGGGCTGACAACGCACTACCACTAAACTCAAACACAACAACAGCAACCACCGCGGCTCAATGGGACTG<br>GAAACGCCACGCGTTGACAGTCCCTCTTGAGGCGTTTGTTATAACTTTCTCGAGGGTATAGCATTCTAG |
| *attC*$_{aadA7}^{bs}$ x *attC*$_{L2}$ | CATTGCGCATTGCGTAGATCTTTTATGTCTAACGCTTGAATTAAGCCGCGCCGCGAAGCGGCGTCGGCTTG<br>AATGAATTGTTAGACATTTTCCATGGCTATATTTCAGCATCATCACATCATCATCATCATCATCACAGGGT<br>AGATCATCATCATCAACACGGGACGACAGCAAATGGCACCCTTCTATAAGCTTTTTGAGTAACCGTTTTGA<br>GTCGAACCCAACGATCCGTTGGTGTTCGAGCTCATCTGTGTTACTCTTTCTCGAGGGTATAGCATTCTAG |

**F.** Staple oligonucleotides for EMSA. Sequences are given in a 5’ to 3’ direction.

| | |
|---|---|
| Staple 1 | CCCTGTGATGATGATGATGATGATGTGATGATGCTGAAATATAGCCATGGACGCAGA<br>TCTACGCAATGCGCAATG |
| Staple 2 | CTAGAATGCTATACCCTCGAGCGCAAGCTTATAGAAGGGTGCCATTTGCTGTCGTCC<br>CGTGTTGATGATGATGATC |

**G.** Plasmids used in this study.

| **Plasmid name** | **Description** | **Reference** |
|---|---|---|
| **Template plasmids for double-*attC* construct** | | |
| double_aadA7_lessCAT_pUC57-Kan-Simple | Containing double-*attC*$_{aadA7}^{bs}$ | This study |
| aadA7-ts_original seq_pUC57-Kan-Simple | Containing double-*attC*$_{aadA7}^{ts}$ | This study |
| aadA7-L2_original seq_pUC57-Kan-Simple | Containing double-*attC*$_{L2}$ | This study |
| VCR-wt_original seq_pUC57-Kan-Simple | Containing double-*attC*$_{VCR2/1}$ | This study |
| VCR-inv_original seq_pUC57-Kan-Simple | Containing double-*attC*$_{VCRinv}$ | This study |

| Plasmids containing MBP-IntI1 | | |
|---|---|---|
| pMAL_c5X_IntI_Y312F_MBP | Catalytically inactive MBP-IntI1$^{Y312F}$ expression plasmid | (*13*) |
| pMAL_c5X_IntI_Y312F_K219A_Y220A_MBP | Catalytically inactive MBP-IntI1$^{A}$ expression plasmid | This study |
| pMAL_c5X_IntI_Y312F_A321X_G322X_MBP | Catalytically inactive MBP-IntI1$^{\Delta C}$ expression plasmid | This study |
| pMAL_c5X_IntI_Y312F_mEGFP_MBP | Catalytically inactive MBP-mEGFP-IntI1$^{Y312F}$ expression plasmid | This study |

**H.** Bacterial strains for *in vivo* conjugation studies.

| Plasmid name | Description | Reference |
|---|---|---|
| **Receptor strain for conjugation** | | |
| V141-V143 | MG1655 Δ*recA* (4826) pL290 | This study |
| V144-V146 | MG1655 Δ*recA* (4826) pL294 | This study |
| V931-V936 | MG1655 Δ*recA* (4826) pV807- pV809 | This study |
| V937-V942 | MG1655 Δ*recA* (4826) pV810- pV813 | This study |
| **Donor strain for conjugation** | | |
| V147-V149 | β2163Δ*km* pV068, pV069 | This study |
| V971-V976 | β2163Δ*km* pV552-pV554 | This study |
| V977-V978 | β2163Δ*km* pV557 | This study |
| W869-W871 | β2163Δ*km* pW654-pW656 | This study |
| W872-W874 | β2163Δ*km* pW657 | This study |
| W863-W865 | β2163Δ*km* p6944 | This study |
| W866-W868 | β2163Δ*km* pV967-pV969 | This study |

**I.** Constructed plasmids for *in vivo* conjugation studies.

| Plasmid name | Description | Notes | Reference |
|---|---|---|---|
| pV068-pV069 | pSW23T*::attC*$_{aadA7}$-*lacIq-attC*$_{aadA7}$ (bs) | *oriV*$_{R6K\gamma}$, *oriT*$_{RP4}$; [Cm$^R$]. The p6944 vector was amplified by PCR with 7169 and 7166 primers. The insert fragment was amplified by PCR from pG291 with 7167 and 7168. Assembly of these two fragments was achieved by performing Gibson Assembly | This study |
| pV552-pV554 | pSW23T*::attC*$_{aadA7}$-*lacIq-attC*$_{aadA7}$ (bs) | *oriV*$_{R6K\gamma}$, *oriT*$_{RP4}$; [Cm$^R$]. EcoRI/SalI digested fragment from pV068 cloned in EcoRI/SalI digested p4747 | This study |
| pV557 | pSW23T*::attC*$_{aadA7}$-*lacIq-attC*$_{aadA7}$ (ts) | *oriV*$_{R6K\gamma}$, *oriT*$_{RP4}$; [Cm$^R$]. EcoRI/SalI digested fragment from pV068 cloned in EcoRI/SalI digested p4746 | This study |
| pV804-pV806 | pSW23T*::attC*$_{aadA7}$-*lacIq-VCR*$_{inv}$ (bs) | Amplification of p6944 by inverse PCR with primers 7237 and 7238 | This study |

| | | | |
|---|---|---|---|
| pV967-pV969 | pSW23T*::attC$_{aadA7}$-lacIq-L2* (bs) | Amplification of p6944 by inverse PCR with o7233 and o7234 | This study |
| pW654-pW656 | pSW23T*::VCR$_{inv}$-lacIq-VCR$_{inv}$* (bs) | Digestion of the pV806 plasmid by EcoRI/BamHI and insertion of the annealed primers o7229/o7230/o7231/o7232 | This study |
| pW657 | pSW23T*::L2-lacIq-L2* (bs) | Digestion of the pV969 plasmid by EcoRI/BamHI and insertion of the annealed primers o7235/o7236 | This study |
| pV807-pV809 | pSC101*::*int1 *mut.* | Amplification of pL294 by inverse PCR with o7246 and o7247 | This study |
| pV810-pV813 | pSC101*::*int1 *trunc.* | Amplification of pL294 by inverse PCR with o7223 and o7245 | This study |

**Table S4. Protein sequences used in this study.** MBP tag is indicated in blue, IntI1$^{Y312F}$ is indicated in yellow, mutated positions are indicated in red.

>MBP-IntI1$^{Y312F}$ (IntI1)

MKIEEGKLVIWINGDKGYNGLAEVGKKFEKDTGIKVTVEHPDKLEEKFPQVAATGDGPDIIFWAHDRFGGYAQSGLLAEIT
PDKAFQDKLYPFTWDAVRYNGKLIAYPIAVEALSLIYNKDLLPNPPKTWEEIPALDKELKAKGKSALMFNLQEPYFTWPLI
AADGGYAFKYENGKYDIKDVGVDNAGAKAGLTFLVDLIKNKHMNADTDYSIAEAAFNKGETAMTINGPWAWSNIDTSKVNY
GVTVLPTFKGQPSKPFVGVLSAGINAASPNKELAKEFLENYLLTDEGLEAVNKDKPLGAVALKSYEEELVKDPRIAATMEN
AQKGEIMPNIPQMSAFWYAVRTAVINAASGRQTVDEALKDAQTNSSSNNNNNNNNNNLGIEGRISHMSMGGRDIGRSENLY
FQGSMKTATAPLPPLRSVKVLDQLRERIRYLHYSLRTEQAYVHWVRAFIRFHGVRHPATLGSSEVEAFLSWLANERKVSVS
THRQALAALLFFYGKVLCTDLPWLQEIGRPRPSRRLPVVLTPDEVVRILGFLEGEHRLFAQLLYGTGMRISEGLQLRVKDL
DFDHGTIIVREGKGSKDRALMLPESLAPSLREQLSRARAWWLKDQAEGRSGVALPDALERKYPRAGHSWPWFWVFAQHTHS
TDPRSGVVRRHHMYDQTFQRAFKRAVEQAGITKPATPHTLRHSFATALLRSGYDIRTVQDLLGHSDVSTTMIFTHVLKVGG
AGVRSPLDALPPLTSER*

>MBP-IntI1$^{Y312F_K219A_Y220A}$ (IntI1$^{A}$)

MKIEEGKLVIWINGDKGYNGLAEVGKKFEKDTGIKVTVEHPDKLEEKFPQVAATGDGPDIIFWAHDRFGGYAQSGLLAEIT
PDKAFQDKLYPFTWDAVRYNGKLIAYPIAVEALSLIYNKDLLPNPPKTWEEIPALDKELKAKGKSALMFNLQEPYFTWPLI
AADGGYAFKYENGKYDIKDVGVDNAGAKAGLTFLVDLIKNKHMNADTDYSIAEAAFNKGETAMTINGPWAWSNIDTSKVNY
GVTVLPTFKGQPSKPFVGVLSAGINAASPNKELAKEFLENYLLTDEGLEAVNKDKPLGAVALKSYEEELVKDPRIAATMEN
AQKGEIMPNIPQMSAFWYAVRTAVINAASGRQTVDEALKDAQTNSSSNNNNNNNNNNLGIEGRISHMSMGGRDIGRSENLY
FQGSMKTATAPLPPLRSVKVLDQLRERIRYLHYSLRTEQAYVHWVRAFIRFHGVRHPATLGSSEVEAFLSWLANERKVSVS
THRQALAALLFFYGKVLCTDLPWLQEIGRPRPSRRLPVVLTPDEVVRILGFLEGEHRLFAQLLYGTGMRISEGLQLRVKDL
DFDHGTIIVREGKGSKDRALMLPESLAPSLREQLSRARAWWLKDQAEGRSGVALPDALERAAPRAGHSWPWFWVFAQHTHS
TDPRSGVVRRHHMYDQTFQRAFKRAVEQAGITKPATPHTLRHSFATALLRSGYDIRTVQDLLGHSDVSTTMIFTHVLKVGG
AGVRSPLDALPPLTSER*

>MBP-IntI1$^{Y312F_A321X_G322X}$ (IntI1$^{\Delta C}$)

MKIEEGKLVIWINGDKGYNGLAEVGKKFEKDTGIKVTVEHPDKLEEKFPQVAATGDGPDIIFWAHDRFGGYAQSGLLAEIT
PDKAFQDKLYPFTWDAVRYNGKLIAYPIAVEALSLIYNKDLLPNPPKTWEEIPALDKELKAKGKSALMFNLQEPYFTWPLI
AADGGYAFKYENGKYDIKDVGVDNAGAKAGLTFLVDLIKNKHMNADTDYSIAEAAFNKGETAMTINGPWAWSNIDTSKVNY
GVTVLPTFKGQPSKPFVGVLSAGINAASPNKELAKEFLENYLLTDEGLEAVNKDKPLGAVALKSYEEELVKDPRIAATMEN
AQKGEIMPNIPQMSAFWYAVRTAVINAASGRQTVDEALKDAQTNSSSNNNNNNNNNNLGIEGRISHMSMGGRDIGRSENLY
FQGSMKTATAPLPPLRSVKVLDQLRERIRYLHYSLRTEQAYVHWVRAFIRFHGVRHPATLGSSEVEAFLSWLANERKVSVS
THRQALAALLFFYGKVLCTDLPWLQEIGRPRPSRRLPVVLTPDEVVRILGFLEGEHRLFAQLLYGTGMRISEGLQLRVKDL
DFDHGTIIVREGKGSKDRALMLPESLAPSLREQLSRARAWWLKDQAEGRSGVALPDALERKYPRAGHSWPWFWVFAQHTHS
TDPRSGVVRRHHMYDQTFQRAFKRAVEQAGITKPATPHTLRHSFATALLRSGYDIRTVQDLLGHSDVSTTMIFTHVLKVGG
**

>MBP-mEGFP-IntI1$^{Y312F}$ (IntI1$_{mEGFP}$)

MKIEEGKLVIWINGDKGYNGLAEVGKKFEKDTGIKVTVEHPDKLEEKFPQVAATGDGPDIIFWAHDRFGGYAQSGLLAEIT
PDKAFQDKLYPFTWDAVRYNGKLIAYPIAVEALSLIYNKDLLPNPPKTWEEIPALDKELKAKGKSALMFNLQEPYFTWPLI
AADGGYAFKYENGKYDIKDVGVDNAGAKAGLTFLVDLIKNKHMNADTDYSIAEAAFNKGETAMTINGPWAWSNIDTSKVNY

GVTVLPTFKGQPSKPFVGVLSAGINAASPNKELAKEFLENYLLTDEGLEAVNKDKPLGAVALKSYEEELVKDPRIAATMEN
AQKGEIMPNIPQMSAFWYAVRTAVINAASGRQTVDEALKDAQTNSSSMVSKGEELFTGVVPILVELDGDVNGHKFSVSGEG
EGDATYGKLTLKFICTTGKLPVPWPTLVTTLTYGVQCFSRYPDHMKQHDFFKSAMPEGYVQERTIFFKDDGNYKTRAEVKF
EGDTLVNRIELKGIDFKEDGNILGHKLEYNYNSHNVYIMADKQKNGIKVNFKIRHNIEDGSVQLADHYQQNTPIGDGPVLL
PDNHYLSTQSKLSKDPNEKRDHMVLLEFVTAAGITLGMDELYKNNNNNNNNNNLGIEGRISHMSMGGRDIGRSENLYFQGS
MKTATAPLPPLRSVKVLDQLRERIRYLHYSLRTEQAYVHWVRAFIRFHGVRHPATLGSSEVEAFLSWLANERKVSVSTHRQ
ALAALLFFYGKVLCTDLPWLQEIGRPRPSRRLPVVLTPDEVVRILGFLEGEHRLFAQLLYGTGMRISEGLQLRVKDLDFDH
GTIIVREGKGSKDRALMLPESLAPSLREQLSRARAWWLKDQAEGRSGVALPDALERKYPRAGHSWPWFWVFAQHTHSTDPR
SGVVRRHHMYDQTFQRAFKRAVEQAGITKPATPHTLRHSFATALLRSGYDIRTVQDLLGHSDVSTTMIFTHVLKVGGAGVR
SPLDALPPLTSER*

**Table S5. Python 3.10.5 packages used in this study.**

| | |
|---|---|
| lumicks.pylake 0.12.1 | pathlib |
| numpy 1.23.1 | Pandas 1.4.3 |
| glob | Scipy 1.8.1 |
| math | Seaborn 0.11.2 |
| statistics | Pingouin 0.5.3 |
| os | Statsmodels 0.13.2 |
| Matplotlib 3.5.2 | Sympy 1.12 |

### Supplementary Protocols

***Double-attC construct preparation***

*dsDNA handle preparation*

PCR amplification of each dsDNA handle was performed in 50 µL volume using 20 ng of template lambda DNA (Promega, USA) using standard Taq polymerase protocol (New England Biolabs, USA) and the designed primer pair (Table S3A) at the annealing temperature of 54°C. The resulting PCR product was cleaned using standard ethanol precipitation protocol.

*Nicking reaction*

Each handle was nicked with a specific restriction enzyme separately according to the manufacturer's instructions (enzymes and buffers from New England Biolabs). The nicking reaction was performed in a thermocycler at 65°C for 1 hour, followed by the inactivation step of 80°C for 20 minutes. The resulting reaction product was cleaned using standard ethanol precipitation protocol.

*dsDNA insert preparation*

The template plasmids (Table S3G) containing the target double-*attC* sequences were commercially acquired (GenScript, USA) and transformed into XL-10 ultracompetent *E. coli* cells using standard heat-shock transformation protocol. The cells were plated on Kanamycin selection plates and grown according to the standard protocol. The resulting culture was used for plasmid DNA extraction using the NucleoSpin plasmid kit (Machery-Nagel, Germany) according to the manufacturer's instructions and eluted with nuclease-free water. PCR amplification of the insert sequence (Table S3E) was performed in 50 µL volume using 100 ng of template plasmid DNA using standard Taq polymerase protocol (New England Biolabs) and the designed primer pair (Table S3B). The resulting PCR product was cleaned using standard ethanol precipitation protocol.

*Selective digestion for single-strand generation*

To remove one strand from the dsDNA insert sequence we used 1 µL (5 units) of lambda exonuclease enzyme (New England Biolabs) on 2.5 µg dsDNA insert. The reaction mix was incubated at 37°C for 30 minutes, followed by the inactivation step of 80°C for 5 minutes.

*Coupling reaction and ligation*

First, one nicked dsDNA handle (5 pmol) is mixed with the ssDNA insert (5 pmol) and left for 15 minutes at room temperature for initial annealing. Then the second nicked dsDNA handle (5 pmol) is added to the mix and the tube is filled until 38 µL with nuclease-free water. The solution is rapidly heated up to 90°C and then slowly cooled down to 16°C for DNA strands annealing. After the coupling step, the reaction solution was supplied with T4 Ligase enzyme and buffer (Jena Bioscience) as well as extra ATP (to the final concentration of 4 mM) and kept at 16°C for 2 hours for ligation. The resulting hybrid DNA molecule was stored at 4°C and used for directly for optical tweezers experiments. All dsDNA products of the assembly were checked on 0.8% agarose gel (1x TBE buffer, running at 90V for 50 minutes) for quality control.

***MBP-IntI1 variants mutagenesis and purification***

*Mutagenesis*

The alanine mutant $IntI1^{A}$ was designed to substitute $Lys^{219}$ and $Tyr^{220}$ with alanines using a QuikChange kit (Agilent). Four nucleotides were substituted in the integrase gene sequence using "primer_fwd_mut_Ala" (Table S3C). The C-terminal truncated mutant $IntI1^{\Delta C}$ was designed to substitute $Ala^{321}$ and $Gly^{322}$ to stop codons using the QuikChange kit (Agilent). Four nucleotides were substituted in the integrase gene sequence using "primer_fwd_mut_stop" (Table S3C).

*Protein expression*

We followed the protein expression and a two-step purification protocol previously developed in the lab (*13*). Each plasmid containing protein variant was transformed into the *E. coli* BL21 (DE3) strain using the heat-shock method and plated on agar containing 50 µg/mL ampicillin for overnight growth at 37°C. One colony was taken from the agar plate to start a 10 mL overnight culture with 50 µg/ml ampicillin at 37°C and 140 rpm. To start a culture 1 L of Terrific broth (TB) medium with 50 µg/mL ampicillin was inoculated using 5 mL of the overnight culture to grow at 37°C. When the culture reached an optical density = 0.6, protein expression was induced by adding isopropyl-D-1-thiogalactopyranoside (IPTG) to a final concentration of 0.3 mM. The induced culture was grown at 14°C overnight. The culture was centrifuged (Beckmann Coulter) at 5500 rpm for 30 min at 4°C, the pellet was collected.

*Protein purification*

Both purification steps were performed on the fast protein liquid chromatography (FPLC) device (ÄKTA Protein Purification System, Cytiva Life Sciences). The first step used the amylose column to separate MBP-tagged protein. However, this purification leaves also free MBP protein that presents as a defined band at 45 kDa on the SDS-PAGE gel. To remove the impurity a second purification step was introduced by using an ion exchange technique. Using difference in theoretical isoelectric points (pI) for the MBP and the MBP-IntI1 variants, separation by charge is possible with the buffer of appropriate pH.

1$^{st}$ step (MBPTrap).

The cell pellet was re-suspended in 60 mL BufferA. Recovered cells were then lysed using EmulsiFlex-C3 (Avestin), and the resulting homogenized cells were centrifuged at 8400 rpm for 1 hour at 4°C. Recovered lysate (supernatant) was filtered using 0.45 µm disposable filters to prevent column clogging. Filtered lysate was loaded onto the ÄKTA FPLC using the superloop at the flow rate of 1 mL/min. The unbound lysate was washed with 5 column volumes of BufferA with a flow rate of 1 mL/min. To elute the bound MBP-tagged protein the gradient from 0-100% of BufferB relative to BufferA was applied and the fractions of 1 mL were collected in a fraction collector.

Collected fractions were analyzed using the sodium dodecyl sulfate-polyacrylamide gel electrophoresis (SDS-PAGE) with 12% acrylamide hand-casted gel (Bio-Rad Minigel system). The gel was run at 150 V for 1 hour and then stained with Comassie by heating it for 30 seconds in the microwave. Afterward, the staining solution was removed and the de-staining solution was added (10% acidic acid) and left overnight on an orbital shaker. After de-staining was complete, the gel was scanned and the bands were analyzed.

2$^{nd}$ step (HiTrap).

Next, to remove the MBP impurity an ion exchange column was used on the same ÄKTA FPLC. The purified fractions were first dialyzed to exchange buffer. Using dialysis cassettes (20000 Da MWCO, Thermo Scientific) and placed in sufficient volume of BufferC on the stirrer at 4$^{o}$C. The buffer was changed after 2 hours and 3 hours and then left overnight. The dialyzed protein was diluted with BufferD (1:4) before loading to lower salt concentration in the sample. Dialysed fraction sample was loaded onto the FPLC using the superloop at the flow rate of 1 mL/min. The unbound lysate was washed with 5 column volumes of BufferD with a flow rate of 1 mL/min. To elute the target protein, the gradient from 0-100% of BufferE relative to BufferD was applied, and the fractions of 1 mL were collected in a fraction collector. Collected fractions were analyzed using SDS-PAGE. High purity fractions were dialyzed to exchange the buffer using the same procedure described above to the final storage BufferF. Protein concentration was calculated using Nanodrop measurements and the known extinction coefficient of the mutant proteins. The samples were aliquoted, frozen with liquid nitrogen, and placed at -70$^{o}$C.

The buffers used were as follows:

• BufferA (running buffer, amylose column): 50mM TRIS-HCl pH7.4, 1M NaCl, 10% glycerol

• BufferB (elution buffer, amylose column): 50mM TRIS-HCl pH 7.4, 1M NaCl, 10mM maltose

• BufferC (4x phosphate buffer exchange): 200mM phosphate buffer pH 7.2, 500mM NaCl, 30% glycerol

• BufferD (running buffer, ion exchange column): 50mM phosphate buffer pH 7.2

• BufferE (elution buffer, ion exchange column): 50mM phosphate buffer pH 7.2, 1M NaCl

• BufferF (protein storage buffer): 50mM TRIS-HCl pH 7.4, 200mM NaCl, 30% glycerol

***EMSA***

We developed a new protocol for non-radioactive EMSA using long ssDNA fragments and bulky fusion proteins that form large molecular complexes and present challenges using standard EMSA conditions. The gel is hand cast using the Bio-Rad Minigel system to produce 5% native polyacrylamide gel (acrylamide/bisacrylamide 37.5:1) with 0.5x TBE and 2.5% glycerol. The running buffer was mixed containing 0.5x TBE and 2.5% glycerol and used to pre-run the gel in the cold (4$^{o}$C) at 80V for 60 minutes.

For optimal integrase-DNA binding conditions, we used the binding buffer (10 mM Tris-HCl, 10 mM NaCl, 40 mM KCl, 1 mM $MgCl_2$, 4 mM EDTA, 1 mM DTT, 0.5 mg/mL BSA, 5% glycerol). The ssDNA of the double-*attC* insert was mixed with the same amount (in pmol) of designed oligonucleotides (Staple 1 and Staple 2) in the final volume of 50 µL and placed in the thermocycler for *attC* hairpin formation and staple annealing (the mix was rapidly heated up to 95°C and then slowly cooled down).

The samples were assembled in 15 µL volume in the following order:

x µL mQ (15 µL – rest)

3 µL binding buffer (5x)

The required amount of IntI protein (diluted from stock with protein storage buffer)

x µL of DNA substrate (2 pmol) – after hairpin formation and staple annealing

The mixture was incubated at room temperature for 15 minutes, then 3 µL of OrangeG dye (6x) was added to the samples to track the run and the mixture was loaded on the gel. The electrophoresis was done in the cold (4°C) at 90 V for 120 minutes. After the run, the gel was placed in a closed box for staining using SYBRGold (Thermo Fisher Scientific) solution (5 µL stock in 50 mL running buffer) and left on the orbital shaker for 30 minutes in the dark. After staining was complete, the gel was rinsed using the running buffer and placed on the plastic film in the imager (Azure 300, Azure Biosystems). The gel was imaged with epi-blue light (LED, 472 nm) and the gel image was analyzed using Fiji.
